# Blood flow regulates *acvrl1* transcription via ligand-dependent Alk1 activity

**DOI:** 10.1101/2024.01.25.576046

**Authors:** Anthony R. Anzell, Amy Biery Kunz, James P. Donovan, Thanhlong G. Tran, Xinyan Lu, Sarah Young, Beth L. Roman

## Abstract

Hereditary hemorrhagic telangiectasia (HHT) is an autosomal dominant disease characterized by the development of arteriovenous malformations (AVMs) that can result in significant morbidity and mortality. HHT is caused primarily by mutations in bone morphogenetic protein receptors *ACVRL1*/ALK1, a signaling receptor, or endoglin (*ENG*), an accessory receptor. Because overexpression of *Acvrl1* prevents AVM development in both *Acvrl1* and *Eng* null mice, enhancing *ACVRL1* expression may be a promising approach to development of targeted therapies for HHT. Therefore, we sought to understand the molecular mechanism of *ACVRL1* regulation. We previously demonstrated in zebrafish embryos that *acvrl1* is predominantly expressed in arterial endothelial cells and that expression requires blood flow. Here, we document that flow dependence exhibits regional heterogeneity and that *acvrl1* expression is rapidly restored after reinitiation of flow. Furthermore, we find that *acvrl1* expression is significantly decreased in mutants that lack the circulating Alk1 ligand, Bmp10, and that BMP10 microinjection into the vasculature in the absence of flow enhances *acvrl1* expression in an Alk1-dependent manner. Using a transgenic *acvrl1:egfp* reporter line, we find that flow and Bmp10 regulate *acvrl1* at the level of transcription. Finally, we observe similar ALK1 ligand-dependent increases in *ACVRL1* in human endothelial cells subjected to shear stress. These data suggest that Bmp10 acts downstream of blood flow to maintain or enhance *acvrl1* expression via a positive feedback mechanism, and that ALK1 activating therapeutics may have dual functionality by increasing both ALK1 signaling flux and *ACVRL1* expression.

## Introduction

Hereditary hemorrhagic telangiectasia (HHT) is an autosomal dominant disorder affecting 1 in 5,000 people worldwide that is characterized by the development of direct connections between arteries and veins, or arteriovenous malformations (AVMs) (reviewed in [1]). Rupture of fragile malformations in the nose, gastrointestinal tract, or brain may cause anemia or hemorrhagic stroke; shunting of blood within lung AVMs may cause brain abscess or embolic stroke; and low systemic vascular resistance caused by liver AVMs may lead to high-output heart failure.

Genes implicated in HHT encode transforming growth factor-β (TGF-β) signaling effectors that function in endothelial cells (ECs). In the TGF-β pathway, ligand binding to a transmembrane receptor complex consisting of two type I and two type II receptors may be facilitated by non-signaling type III receptors. Complex formation allows the type II receptor to phosphorylate the type I receptor, which in turn initiates intracellular signaling by phosphorylation of receptor-specific Smad (R-Smad) proteins. R-Smads form a heterotrimer with Smad4, translocate to the nucleus, and regulate transcription of target genes. Mutations in *ENG*, encoding the type III receptor endoglin, result in HHT1 [2] while mutations in *ACVRL1*, encoding the type I receptor, activin A receptor-like type 1 (ALK1), result in HHT2 [3]. Mutations in these two genes account for up to 96% of clinically diagnosed HHT [4].

The only medical therapies available to HHT patients block angiogenesis or speed clotting. These agents are not effective in all patients and carry risk of increased thrombosis [5,6]. Accordingly, there is a critical unmet need for development of targeted HHT therapeutics. Given that HHT is an autosomal dominant disorder caused largely by loss-of-function variants [7–10], haploinsufficiency may be the genetic mechanism of disease, and there is some evidence to support this assumption [9,11,12].

However, there is compelling evidence for mutation of the wild type allele of the germline-mutated gene in at least some HHT telangiectasias, supporting the alternative hypothesis that mosaic loss of function caused by somatic second hits seeds HHT lesions [13]. Regardless of genetic mechanism, restoring ALK1 signaling by inducing *ACVRL1* expression may be a practical approach to therapy. If haploinsufficiency causes disease, this approach would benefit HHT2 (*ACVRL1*) and HHT1 (*ENG*) patients, as *ENG* enhances but is not required for ALK1 signaling [10], and overexpression of *Acvrl1* prevents AVM development in *Eng* null mice [14]. If somatic second hits cause diseases, then this approach would still benefit HHT1 (*ENG*) patients with intact *ACVRL1*, who compose approximately half of the HHT patient population.

There is limited information regarding the regulatory elements that govern *ACVRL1* expression. The human *ACVRL1* promoter contains validated Sp1 and Kruppel-like factor 6 (KLF6) binding sites, which are required for basal and/or injury-induced *ACVRL1* transcription in ECs [15,16], and UBR5, a nuclear E3 ubiquitin ligase, may be involved in negative regulation by antagonizing Sp1 binding [17]. In addition, functional binding sites for the ETS family transcription factors (ETS1, FLI1) and FOXF1 have been identified in the mouse *Acvrl1* promoter, and a 2.2 kb element in mouse *Acvrl1* intron 2 (which is equivalent to human and zebrafish intron 1) is required for arterial EC-localized *Acvrl1* expression [18–21]. However, how this intronic fragment directs arterial expression is unknown.

Several lines of evidence suggest that *ACVRL1* expression is regulated by blood flow. In zebrafish, *acvrl1* is expressed predominantly in arterial ECs proximal to the heart, and *acvrl1* expression depends on blood flow [22]. However, whether the control mechanism involves transcriptional or post-transcriptional regulation has not been investigated, and whether mechanical force imparted by blood flow or circulating factors carried by blood flow explain this requirement is unclear. Here, we demonstrate in zebrafish that blood flow enhances or maintains *acvrl1* in a Bmp10-dependent manner and uncover regional heterogeneity in *acvrl1* flow dependence, with arterial EC sensitivity to flow and *bmp10* increasing with distance from the heart. Moreover, we find that in the absence of blood flow Bmp10/Alk1 signaling is sufficient to drive *acvrl1* expression and that this process is regulated at the level of transcription, at least in part via an enhancer element in intron 1. Finally, we observe ALK1-ligand dependent increases in *ACVRL1* expression in human ECs subjected to simulated blood flow. Together, these data suggest that the blood flow requirement for *acvrl1* expression stems primarily from the requirement for circulating Bmp10, and that protein ligands or synthetic ALK1 agonists may enhance *ACVRL1* expression and ALK1 signaling and serve as viable therapeutics for HHT patients.

## Materials and Methods

### Zebrafish lines and maintenance

Zebrafish (*Danio rerio*) were maintained according to standard protocols [23]. To maintain optical clarity of embryos, we added 0.003% 1-phenyl-2-thiourea (PTU; Sigma, St. Louis, MO, USA) to 15% Danieau embryo medium [8.5 mM NaCl, 1 mM KCl, 0.06 mM MgSO_4_, 0.9 mM Ca(NO_3_)_2_, 0.75 mM HEPES] as early as 8 hours post-fertilization (hpf). Mutant lines *acvrl1^y6^* (p.L249F), *bmp10^pt527^* (p.T45Rfs*4), and *bmp10-like^sa11654^* (p.R180*) were previously described [24,25]. The *Tg*(*fli1a.ep*:*mRFP*)*^pt505^*line drives membrane-RFP expression under an endothelial-specific enhancer and has been described [22].

To generate *Tg(acvrl1e5:egfp),* we used Gateway cloning (Thermo Fisher Scientific, Waltham, MA, USA) to assemble a DNA construct flanked by *Tol2* transposon arms [26,27] and containing a 1910-bp fragment from the zebrafish *acvrl1* gene (GRCz11 23:27854594-27856491; final 3 bases of exon 1 plus 1907 bases of 5’ intron 1) upstream of a basal promoter derived from carp β-actin and coding sequence for EGFP. We injected 25 pg *acvrl1e5:egfp* DNA plus 25 pg tol2 transposase synthetic mRNA (mMessage mMachine Sp6, Thermo Fisher Scientific) into one-cell stage embryos to establish *Tg*(*acvrl1e5:egfp*)^pt517^. To map the insertion site, we digested genomic DNA from a *Tg(acvrl1e5:egfp)^pt517^* fish with HaeIII, ligated digestion products, performed inverse PCR using primers specific to the 3’ end of the *acvrl1e5:egfp* plasmid (5’-CAGTAATCAAGTAAAATTACTC-3’; 5’-AATGCACAGCACCTTGACCTGG-3’), and sequenced the PCR product. We confirmed insertion in chromosome 20 by PCR and sequencing. For the 5’ insertion site, we used primers 5’-GACGCAGCATTTGCAGAGCTAAACTA-3’ (*pinx1* intron 6) and 5’-AGTACAATTTTAATGGAGTACT-3’ (plasmid backbone) to amplify a 197 bp product. For the 3’ insertion site, we use primers 5’-CTCAAGTAAAGTAAAAATCC-3’ (plasmid backbone) and 5’-GAGACACTCAACATAGGCTCTAGG-3’ (*pinx1* intron 6) to amplify a 1191 bp product.

To generate *Tg(acvrl1e5:egfp)^pt552^*, we used Crispor [28] to design Cas9 guide RNAs (gRNAs) to delete a 5’ region of the *Tg*(*acvrl1e5:egfp*)^pt517^ transgene. The 5’ target (PAM sequence underlined), CCAGATGGGCCTCTAGC<u>GTT</u>, was located in the plasmid backbone of the transgene, and the 3’ target, GTGCTTACGAATACACTG<u>GTT</u>, was located in *acvrl1* intron 1. We generated single guide RNAs (sgRNAs) [29] from PCR templates using MEGAshortscript (Thermo Fisher Scientific), co-injected one-cell stage embryos with RNPs containing 25 pg of gRNA and 500 pg of NLS-Cas9 protein (PNA Bio, Thousand Oaks, CA USA), and raised these fish to adulthood. We assayed deletion in embryos from outcrosses using PCR (primers 5’-CAATCCTGCAGTGCTGAAAAGCCTC-3’ and 5’-TCAGCTGCTCACAATAGCCT-3’; 1615 bp product in *Tg*(*acvrl1e5:egfp*)^pt517^). Sanger sequencing confirmed a 1242 bp deletion in stable line *Tg(acvrl1e5:egfp)^pt552^*.

### Morpholinos and drug exposures

To prevent initiation of heartbeat, we injected 4 ng of a translation blocking *tnnt2a* morpholino-modified antisense oligonucleotide (5′-CATGTTTCGTCTGATCTGACACGCA-3’ [30]) or standard control morpholino (5’-CCTCTTACCTCAGTTACAATTTATA-3’) (GeneTools, Philomath, OR, USA) into embryos at the one-to two-cell stage. To transiently stop heartbeat, we incubated embryos in 800 μg/mL tricaine methanesulfonate (Syndel, Ferndale, Washington, USA) in 15% Danieau/0.003% PTU for indicated times.

### Microinjection of BMP10 into the vasculature

We performed microinjection into the caudal division of the internal carotid artery (CaDI) as previously described [31] with minor modifications. To determine whether BMP10 can serve as a proxy for blood flow in modifying *acvrl1* expression, we treated wild type or *Tg(acvrl1e5:egfp)^pt552^* embryos with 1.6 mg/mL tricaine at 32 hpf or 44 hpf, respectively, for 1-2 minutes. Once heartbeat ceased, we immediately embedded embryos in 0.75% NuSieve GTG low-melt agarose (Lonza, Rockland, ME, USA) containing 800 μg/mL tricaine and microinjected 2 nL of 3.33 µM recombinant human BMP10 growth factor dimer (generated by Dr. Andrew Hinck, University of Pittsburgh, PA, USA) or vehicle (0.1 M KCl, 0.5% phenol red) into the left CaDI. Following microinjection, we unembedded embryos and transferred them to 15% Danieau/0.003% PTU/800 μg/mL tricaine for 4 hours to maintain a no-flow state, then fixed embryos in 4% paraformaldehyde for in situ hybridization. Data represent 4 independent experiments.

To determine whether BMP-mediated *acvrl1* expression depends on Alk1 activity, we performed a similar experiment in *tnnt2a* morpholino- or control morpholino-injected embryos from an *acvrl1^y6/^*^+^ incross. At 32 hpf, we embedded embryos in 0.75% NuSieve GTG low-melt agarose/800 μg/mL tricaine and injected human BMP10 into the CaDI as described above. After 4 hours of recovery in 15% Danieau/0.003% PTU, we fixed embryos for situ hybridization. Data represent 5 independent experiments.

### In situ hybridization

We performed whole-mount in situ hybridization as previously described [24] using digoxigenin-labeled riboprobes, anti-digoxigenin-AP F(ab) (Sigma-Aldrich), and NBT/BCIP substrate (Sigma-Aldrich). Riboprobes complementary to *acvrl1*, *cdh5*, *egfp*, *sox7*, and *pinx1* were generated from pCRII-TOPO vectors or PCR products using a digoxigenin RNA labeling kit (Sigma-Aldrich). For Supplementary Fig. 3, we use a single *acvrl1* riboprobe targeting exon 9-11, including part of the 3’ UTR, using a PCR product generated from cDNA using primers 5’-CGATACATGGCACCAGAGGT-3’ and 5’-<u>ATTTAGGTGACACTATAGAAGTG</u>CCCACAGAGAACGAATGTCA-3’ (SP6 site underlined) [24]. For all other figures, to increase the sensitivity of *acvrl1* detection, we used a cocktail of three nonoverlapping *acvrl1* riboprobes targeting: i) exon 1-5, primers 5’-TTGCCGCCCGTTATGAGAATAC-3’ and 5’-<u>ATTTAGGTGACACTATAGAAGTG</u>ACAACACCAGCAACGGCACTC-3’; ii) exon 7-9, primers 5’-AAAGGCCGTTATGGTGAGGTATG-3’ and 5’-<u>ATTTAGGTGACACTATAGAAGTG</u>GGTGCCCACTCTCGGATTG-3’; and iii) exon 9-11, as described above. We allowed the color reaction to proceed for 1 day at room temperature and fixed all embryos stained with a given probe at the same time. We captured bright-field images using an MVX-10 MacroView macro zoom microscope equipped with an MV PLAPO 1x/0.25 NA objective, 2x magnification changer, and DP71 camera with cellSens Entry 1.16 software, build 15404 (Olympus America, Center Valley, PA, USA). We coded samples before blindly scoring qualitative expression as “positive” or “negative” in individual *acvrl1-*positive arterial vessels: namely, the first aortic arch (AA1), internal carotid artery (ICA), caudal division of the internal carotid artery (CaDI), basal communicating artery (BCA), and lateral dorsal aorta (LDA). When relevant, after imaging and phenotype scoring, we genotyped embryos using established assays for *acvrl1^y6^* [22], *bmp10^pt527^*, and *bmp10-like^sa11654^* [25]. For Supplementary Fig. 1, we allowed the color reaction to proceed for 5 days at room temperature, embedded embryos in 4% NuSieve GTG agarose (Lonza, Rockland, ME, USA), and sectioned at 50 µm (*acvrl1*) or 100 µm (*cdh5*) with a VT1000S vibratome (Leica Microsystems, Buffalo Grove, IL, USA). We mounted sections with Vectashield mounting media and captured brightfield images using an Olympus BX51 upright compound microscope outfitted with a UPAN FLN 20X/0.5 NA dry objective and DP71 camera as described above. All images were compiled with Adobe Photoshop 9.0.2 or CS2 24.1.1 and Adobe Illustrator V24.1 (Adobe Systems, San Jose, CA, USA).

### Cell culture

We cultured pooled human umbilical vein endothelial cells (HUVEC; C-12203, Promocell, Heidelberg, Germany) at 37°C, 5% CO_2_ in endothelial growth medium 2 (EGM2, consisting of endothelial basal medium-2 supplemented with growth factors; Promocell) and 1X antibiotic-antimycotic (Thermo Fisher Scientific, USA). For standard growth conditions, we used all EGM2 supplements, including 2% fetal bovine serum (FBS). For serum starvation conditions, we included all EGM2 supplements except FBS (0% FBS). HUVECs were at passage 6 (P6) for all experiments.

### Mesofluidic channel fabrication and experimental setup

We designed mesofluidic channels (13 mm wide x 200 μm high x 54 mm long), modeled negative geometries using SolidWorks 2016 (Dassault Systemes, Velizy-Vallicoublay, France), built 3D-printed negative molds using Accura 10 polymer and a VIPER Si2 stereolithography system (3D Systems, Rock Hill, South Carolina, USA), and used replica molding to fabricate channels out of polydimethylsiloxane (Sylgard 184 PDMS; Dow Corning, Midland, MI, USA), as we described previously [32]. We prepared cured molds and bonded them to a 24 mm x 60 mm #1.5 pre-cleaned coverslip as previously described [32]. To simulate flow, we generated a sterile circuit by pumping 50 mL of medium from a 50 mL conical tube reservoir using a Masterflex Ismatec Reglo Independent Channel Control Peristaltic Pump (MFLX78001-80, 4-channel, 8-roller; Avantor/VWR, Radnor, PA, USA) fitted with 1.52 mm inner diameter 3-stop tubing (95625-36, PVC pump tubes, Ismatec). Media then passed through a high flow bubble trap (06BT-HF, Diba Omnifit PEEK with 10 µm PTFE filter, Cole-Parmer, Vernon Hills, IL, USA) and finally into the mesofluidic device using tygon tubing (95702-00, 0.031” inner diameter, 0.094” outer diameter; Cole Parmer). The entire flow setup, including the pump, was placed in 37°C, 5% CO_2_ for the duration of the experiment.

### *In vitro* flow simulation

We treated confluent P5 HUVECs with 0.05% trypsin-EDTA (Thermo Fisher Scientific) for 3 minutes at 37°C, centrifuged at 1000 x g for 5 minutes, and resuspended in EGM2 with 2% FBS. We diluted HUVECs to 2.16 million cells/mL and seeded 200 μl of suspension into mesofluidic channels pre-coated with 50 µg/mL fibronectin (Sigma-Aldrich). For experiments with varying shear stress magnitude, we incubated HUVECs in the mesochannels under static conditions for a 16-hour acclimation period, followed by 28 hours of flow. For experiments testing the role of ALK1 ligands in flow-induced *ACVRL1* expression, we incubated HUVECs in mesochannels under static conditions for a 16-hour acclimation period, followed by a 4-hour serum starvation period (0% FBS), then 28 hours of flow. To generate ALK1 ligand-depleted EGM2, we pre-treated 1 mL of FBS with 2.5 mg of recombinant human ALK1-Fc (R&D systems) and rocked at room temperature for 2 hours before adding to 49 mL EGM2 for a final concentration of 50 µg/mL ALK1-Fc. For all experiments, following 28 hours of flow, we washed HUVECs once with Dulbecco’s PBS, harvested with 200 μL of RLT buffer, and isolated total RNA using RNeasy mini kit (Qiagen, Germantown, MD, USA). We generated cDNA from 1 µg of RNA using the SuperScript IV First Stand Synthesis kit (Thermo Fisher Scientific) and analyzed by RT-qPCR using Quant Studio 12K Flex Real Time PCR system version 1.1.2 (Applied Biosystems by Life Technologies, Thermo Fisher Scientific) and validated (efficiency > 1.8) TaqMan assays (Thermo Fisher Scientific) for *ACVRL1* (Hs00953798_m1), *ID1* (Hs03676575_s1), *KLF2* (Hs00360439_g1), and *GAPDH* (Hs02786624_g1). We analyzed data using the ΔΔCt method [33]. For experiments testing shear stress magnitude, data represent 3 independent experiments. For experiments testing the role of ALK1 ligands at 8 and 12 dyn/cm^2^, data represent 4 and 3 independent experiments, respectively.

### Confocal and two-photon imaging

We performed live confocal and two-photon imaging of *Tg(acvrl1e5:egfp)^pt517^;Tg(fliep:mRFP-CAAX)^pt505^*zebrafish embryos as described previously [34]. We anesthetized embryos in 160 μg/mL tricaine and embedded them in 5% methyl cellulose (Sigma-Aldrich) or 1% NuSieve GTG low-melt agarose. We captured images using an upright TCS SP5 confocal microscope (Leica Microsystems, Wetzlar, Germany) outfitted with an HCX APO L 20x/0.95 or HCX IRAPO L 25x/0.95 water-dipping objective, a 561 nm diode (for RFP), a Mai Tai DeepSeeTi Sapphire laser (900 nm for EGFP; Newport/Spectra Physics, Santa Clara, CA, USA), a galvo scanner (600 Hz), and HyD spectral and nondescanned detectors. We collected Z-series (1.48 μm steps) using 3x line averaging and 1.7x zoom. We generated maximum intensity projections in both channels using LAS AF (version 3.0.0 build 834) and generated figures using Adobe Photoshop.

### Statistics

Qualitative expression levels of *acvrl1* comparing the interaction of flow and Bmp10 were compared by two-tailed Fisher’s exact test using SPSS (Version 28, IBM Inc., Chicago, IL, USA) [35]. All other experiments analyzing qualitative expression levels of *acvrl1* and *egfp* were compared by two-tailed Fisher’s exact test using the MedCalc Software Fisher’s exact probability calculator (MedCalc Software Ltd; https://www.medcalc.org/calc/fisher.php, Version 22.009). All HUVEC data were analyzed using Prism 9.5.1 (GraphPad, Boston, MA, USA). HUVEC data comparing different shear stress magnitudes were analyzed by one-way ANOVA with Tukey’s post-hoc analysis for comparisons among all groups. HUVEC data comparing the effects of ALK1-Fc treatment and shear stress were analyzed by two-way ANOVA with Tukey’s post-hoc analysis to detect interactions and differences among groups.

Differences were considered significant at *p* < 0.05.

## Results

### *acvrl1* expression in arterial ECs is sensitive to lack of blood flow, with flow dependence increasing with distance from the heart

In the zebrafish embryo, *acvrl1* expression is first detectable after the onset of blood flow (∼24-26 hpf) in arterial endothelial cells, with relatively high expression in cranial arteries most proximal to the heart [22]. Blood leaves the heart through the first aortic arch (AA1) and flows cranially into the bilateral internal carotid arteries (ICA), the caudal divisions of the ICAs (CaDI), and into the basal communicating artery (BCA), as well as caudally into the paired lateral dorsal aortae (LDA) and single dorsal aorta (DA). All of these arteries, as well as the optic artery (OA), express relatively high levels of *acvrl1*. From the BCA, blood then flows through posterior communicating segments (PCS) and into the basilar artery (BA); these arteries also express *acvrl1*, but at relatively low levels (Supplementary Fig. 1). Other arterial ECs (for example, central arteries in the brain, intersegmental arteries in the trunk) and all venous ECs are *acvrl1-*negative by in situ hybridization. How this striking demarcation of expression domains is established and maintained is unknown.

We previously reported that *acvrl1* expression requires blood flow: in zebrafish *tnnt2a* mutant embryos, which never initiate heartbeat, we could not detect *acvrl1* at 36 hpf by in situ hybridization when using a single riboprobe complementary to exons 9-11 [22]. Because oxygen needs of the embryo at this time can be met by diffusion [36], we concluded that this lack of expression does not represent a hypoxic response, and that *acvrl1* expression must require mechanical force imparted by and/or circulating factors distributed by blood flow. However, comparing *tnnt2a* morphants to control morphants and using a more sensitive protocol that employs a cocktail of three non-overlapping riboprobes, we now report spatiotemporal variation in the flow-dependence of *acvrl1* (Fig. 1A-D; statistical analysis in Supplementary Table 1). In control embryos, *acvrl1* was detected in nearly all scored vessels at 28, 32, and 36 hpf (Fig. 1B-D). Comparing *tnnt2a* morphants (no blood flow) to controls, the intensity of the *acvrl1* hybridization signal was subjectively decreased in AA1 and the ICA, the arteries most proximal to the heart; however, *acvrl1* remained detectable in *tnnt2a* morphants in AA1 (100%) and the ICA (80-90%) (Fig. 1B-D). By contrast, in the CaDI and BCA, which are more distal cranial vessels, only 15-20% of *tnnt2a* morphants expressed *acvrl1* at 28 or 32 hpf, with no detectable expression at 36 hpf (Fig. 1B-D). A similar spatiotemporal trend in flow sensitivity was observed in the LDA and DA in the trunk. In *tnnt2a* morphants, *acvrl1* expression in the proximal LDA was detected in ∼70% of embryos at 28 hpf, declining to ∼8% of embryos at 36 hpf, and in the more distal DA, in ∼48% of embryos at 28 hpf, declining to ∼3% of embryos at 36 hpf (Fig. 1B-D). Notably, the pan-endothelial marker, *cdh5*, was unaffected in *tnnt2a* morphants compared to controls (Supplementary Fig. 2A). Collectively, these results demonstrate that blood flow is required to maintain and/or enhance but not initiate *acvrl1* expression, and that *acvrl1* dependence on blood flow increases in a proximal-to-distal fashion with respect to the heart.

**Fig. 1.**
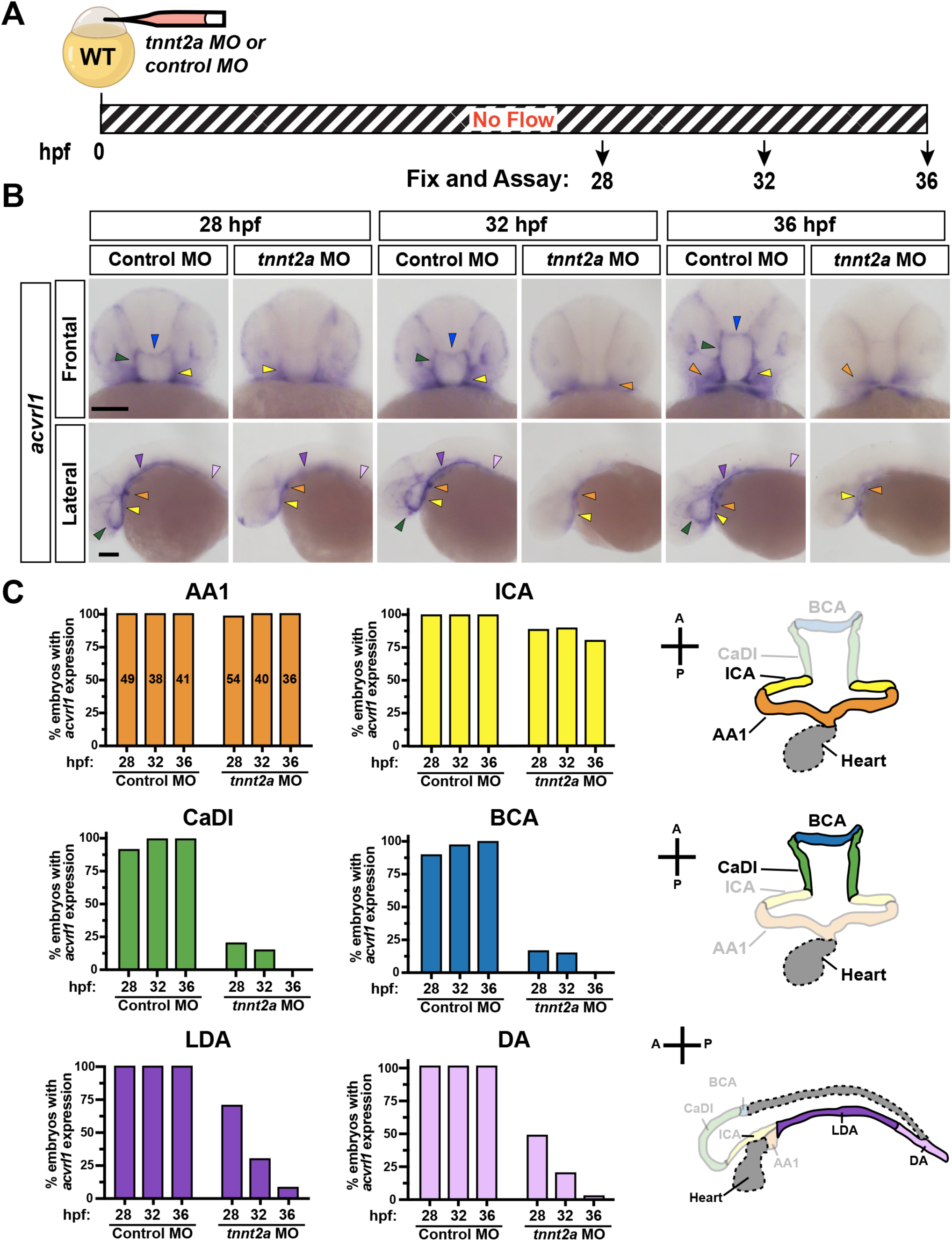
*acvrl1* expression is maintained or enhanced by blood flow**. a** Experimental design. Embryos were injected at the 1-cell stage with *tnnt2a* or control morpholino, fixed at indicated timepoints, and assayed by *in situ* hybridization for *acvrl1*. **b** Representative *acvrl1* images, frontal (top) and lateral (bottom) views. Color-coded arrows mark presence of signal in indicated vessels. Scale bars = 100 µm. **c** Percent of embryos with *acvrl1* expression detected in indicated vessels in control and *tnnt2a* morphants at indicated times. The number of embryos analyzed per group, summed across 3 independent experiments, is displayed in AA1 graph. At right, schematics of vessels shown in frontal (top, middle) and lateral (bottom) views. Differences between groups for each vessel were computed by Fisher’s exact test and are reported in Supplementary Table 1. Color coding in **b**,**c**: first aortic arch (AA1, orange), internal carotid artery (ICA, yellow), caudal division of the ICA (CaDI, green), basal communicating artery (BCA, blue), lateral dorsal aorta (LDA, dark purple), dorsal aorta (DA, light purple)

### *acvrl1* expression in arterial ECs responds to both cessation and restoration of blood flow, with flow dependence increasing with distance from the heart

Having previously demonstrated that cessation of blood flow for 8 hours (32-40 hpf) results in loss of *acvrl1* expression [22], we next asked whether *acvrl1* might recover if blood flow is allowed to resume. To this end (Supplementary Fig. 3A), we treated wild type embryos with 800 μg/mL tricaine at 32 hpf to stop heartbeat and collected embryos at 34 hpf (2 hours without flow) and 40 hpf (8 hours without flow). We then washed embryos out of tricaine at 40 hpf and collected embryos at 42 hpf (2 hours recovery) and 48 hpf (8 hours recovery). Heartbeat in tricaine-treated embryos was noticeably slowed by 15 minutes, and red blood cell circulation ceased by 30-60 minutes post-treatment (32.5-33 hpf). After washout at 40 hpf, heartbeat was detectable by 15-30 minutes and blood flow was indistinguishable from sibling controls within 1-to-2 hours (41-42 hpf).

At all time points, we detected *acvrl1* expression in all expected arterial vessels in most sibling control embryos; the lack of expression in some control embryos likely reflects the low sensitivity of the single-probe *acvrl1* in situ hybridization that was used for these experiments (Supplementary Fig. 3B,C). Tricaine exposure for as short as 2 hours (34 hpf) decreased *acvrl1* expression in all assayed vessels, with effects more pronounced in distal (CaDI, BCA) versus proximal (AA1, ICA, LDA) segments. With 8 hours of tricaine exposure (40 hpf), *acvrl1* expression was undetectable in nearly all embryos, as we have previously reported [22], with just ∼5% of embryos retaining some expression in AA1 only. Remarkably, within 2 hours after tricaine removal (42 hpf) and subsequent reinitiation of heartbeat and blood flow, *acvrl1* expression was restored in most arterial vessel segments, with more complete restoration in proximal (AA1, ICA, LDA) versus distal (CaDI, BCA) segments. Within 8 hours of tricaine removal (48 hpf), *acvrl1* expression was fully restored. Notably, *cdh5* expression remained unchanged in tricaine-treated versus control embryos at all times (Supplementary Fig. 3D). Together, these data reveal the rapid dynamics of *acvrl1* expression changes in response to loss and restoration of blood flow and confirm that *acvrl1* expression is more dependent on blood flow in distal versus proximal arterial ECs.

### Bmp10 is required for the enhancement and/or maintenance of *acvrl1* expression

Blood flow imparts mechanical force on the vessel wall and also carries a plethora of circulating factors, including ALK1 ligands BMP9 and BMP10 [37]. We previously demonstrated that *acvrl1* mRNA is downregulated in *acvrl1^y6/y6^*zebrafish embryos that harbor a missense mutation that generates a kinase-dead protein [22,24]. This missense mutation is unlikely to confer transcript instability. Therefore, we reasoned that Alk1 activity may be required in a positive feedback mechanism to drive *acvrl1* mRNA expression, and that effects of flow cessation on *acvrl1* expression may be caused by loss of circulating Alk1 ligand and consequent loss of Alk1 activity. In zebrafish, there are two *BMP10* paralogs, *bmp10* and *bmp10-like*, and *bmp10;bmp10-like* double mutants fully phenocopy *acvrl1* mutants, developing embryonic lethal cranial AVMs. By contrast, *bmp9* mutants develop no phenotype [25]. Accordingly, Bmp10 is the only required Alk1 ligand in embryonic zebrafish. Therefore, to determine whether Alk1 ligands are required for *acvrl1* expression, we assayed expression in embryos derived from *bmp10^pt527/+^;bmp10-like^sa11654/sa11654^* incross at 28, 32, and 36 hpf (Fig. 2A-C; statistical analysis in Supplementary Table 2). Embryos were scored blinded and genotyped after scoring; we present data from *bmp10^+/+^;bmp10-like^sa11654/sa11654^* sibling controls (no embryonic phenotype) and *bmp10^pt527/pt527^;bmp10-like^sa11654/sa11654^* double mutants (AVMs after 36 hpf). In the *bmp10;bmp10-like* double mutants, although the intensity of the *acvrl1* hybridization signal was subjectively decreased in AA1 and ICA compared to controls, it was detectable in AA1 (100%) and the ICA (∼73-94%) at 28-36 hpf (Fig. 2B-C). The CaDI and BCA were more sensitive to the loss of Bmp10, with ∼42-50% of *bmp10;bmp10-like* double mutants expressing *acvrl1* here at 28 hpf and progressive loss of signal over time. Finally, the LDA and DA were generally refractory to Bmp10 loss at 28 hpf but exhibited progressive loss of signal at 32 and 36 hpf (Fig. 2B-C). No differences were observed in *cdh5* in *bmp10^pt527^;bmp10-like^sa11654^* double mutants compared to controls at any time (Supplementary Fig. 2B) Together, these data reveal that *acvrl1* expression exhibits similar regional heterogeneity in sensitivity to the loss of blood flow or the loss of Bmp10, with proximal vessels (AA1, ICA) marginally dependent on both to enhance *acvrl1* expression, and distal cranial vessels (CaDI, BCA) and trunk vessels (LDA, DA) strongly dependent on both to maintain *acvrl1* expression.

**Fig. 2.**
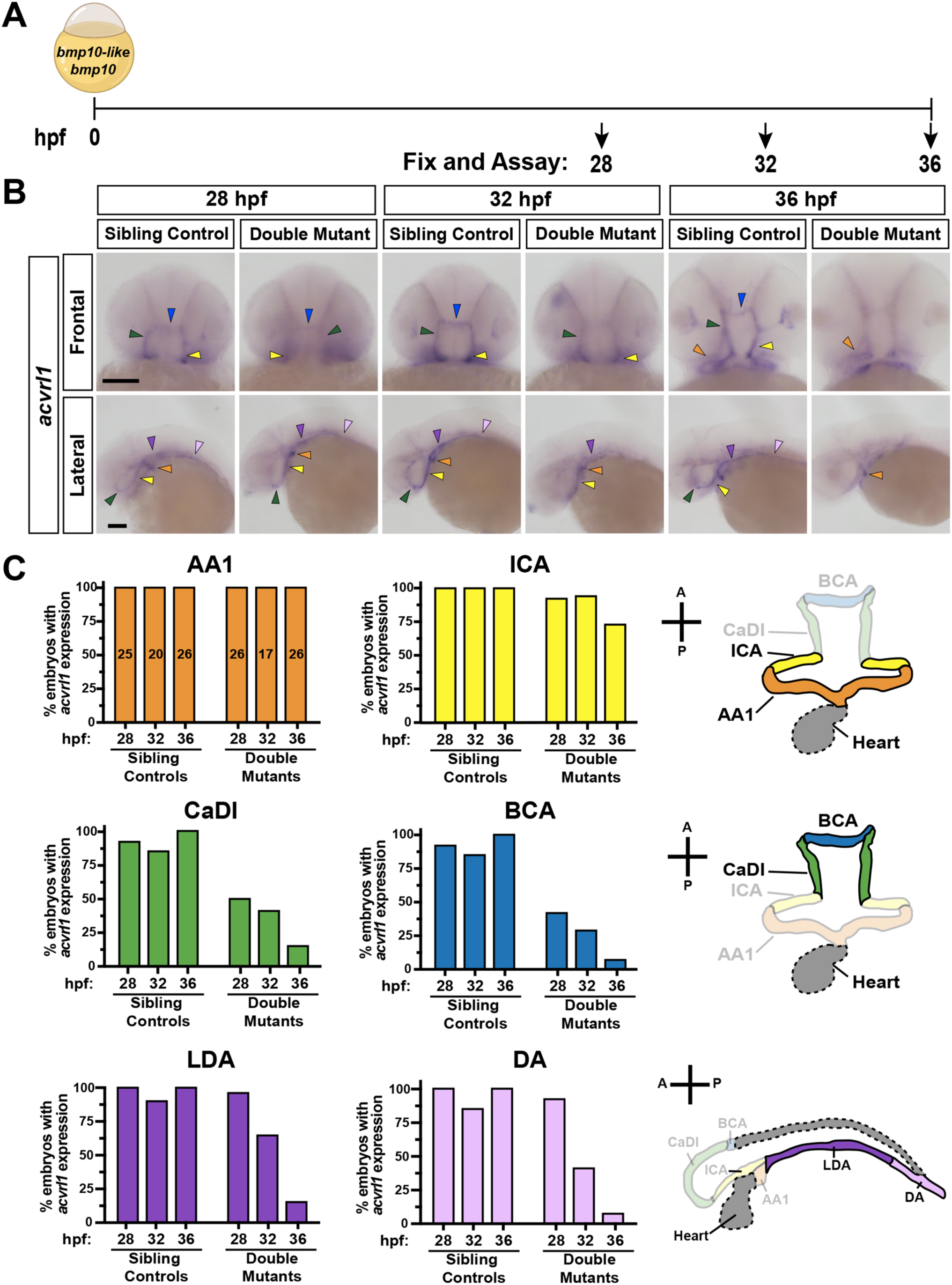
*acvrl1* expression is maintained or enhanced by Bmp10**. a** Experimental design. Embryos generated from a *bmp10^pt527/+^;bmp10-like^sa11654/sa11654^*incross were fixed at the indicated timepoints and assayed by *in situ* hybridization for *acvrl1*. **b** Representative *acvrl1* images, frontal (top) and lateral (bottom) views. Color-coded arrows mark presence of signal in indicated vessels. Scale bar = 100 µm. **c** Percent of embryos with *acvrl1* expression detected in indicated vessels in sibling controls (*bmp10^+/+^;bmp10-like^-/-^*) and double mutants (*bmp10^-/-^;bmp10-like^-/-^*). The number of embryos analyzed per group, summed across 3 independent experiments, is displayed in AA1 graph. Differences between groups for each vessel were computed by Fisher’s exact test and are reported in Supplementary Table 2. Color coding in **b**,**c**: first aortic arch (AA1, orange), internal carotid artery (ICA, yellow), caudal division of the ICA (CaDI, green), basal communicating artery (BCA, blue), lateral dorsal aorta (LDA, dark purple), dorsal aorta (DA, light purple)

### Bmp10 is required for restoration of *acvrl1* expression upon resumption of blood flow

Thus far, our data demonstrate that *acvrl1* expression is dependent on blood flow and the circulating Alk1 ligand, Bmp10, and that *acvrl1* expression decreases with cessation of blood flow and is quickly restored when blood flow resumes. To examine the interaction between blood flow and Bmp10, we stopped heartbeat by treating embryos derived from *bmp10^pt527/+^;bmp10-like^sa11654/sa11654^* incrosses with tricaine from 32-36 hpf, washed embryos out of tricaine at 36 hpf, and allowed heartbeat and blood flow to recover until 40 hpf (Fig. 3A). We assayed *acvrl1* expression at 36 and 40 hpf in *bmp10^+/+^;bmp10-like^sa11654/sa11654^*sibling controls and *bmp10^pt527/pt527^;bmp10-like^sa11654/sa11654^* double mutants, and focused our analysis on the CaDI, BCA, LDA, and DA (Fig. 3B-C; statistical analysis in Supplementary Table 3) because *acvrl1* expression in these vessels is highly dependent on both blood flow and Bmp10. As expected, in sibling controls, *acvrl1* expression was lost in each of these vessels after 4 hours of stopped flow (36 hpf, tricaine) and was restored to normal levels after 4 hours of resumed flow (40 hpf, tricaine). Also as expected, in *bmp10;bmp10-like* double mutants, *acvrl1* expression was absent in these vessels in most embryos at 36 hpf, regardless of flow state. Notably, after 4 hours of resumed flow (40 hpf), *acvrl1* expression was not restored in any vessels in *bmp10;bmp10-like* double mutants. No differences were observed in *cdh5* expression in any condition (Supplementary Fig. 4). These data suggest that the requirement for blood flow may not reflect a role for hemodynamic force in maintenance of *acvrl1* expression. Instead, the role of blood flow in maintenance of *acvrl1* expression may reflect the need to distribute circulating Bmp10 to induce Alk1 activity, which in turn induces *acvrl1* expression.

**Fig. 3.**
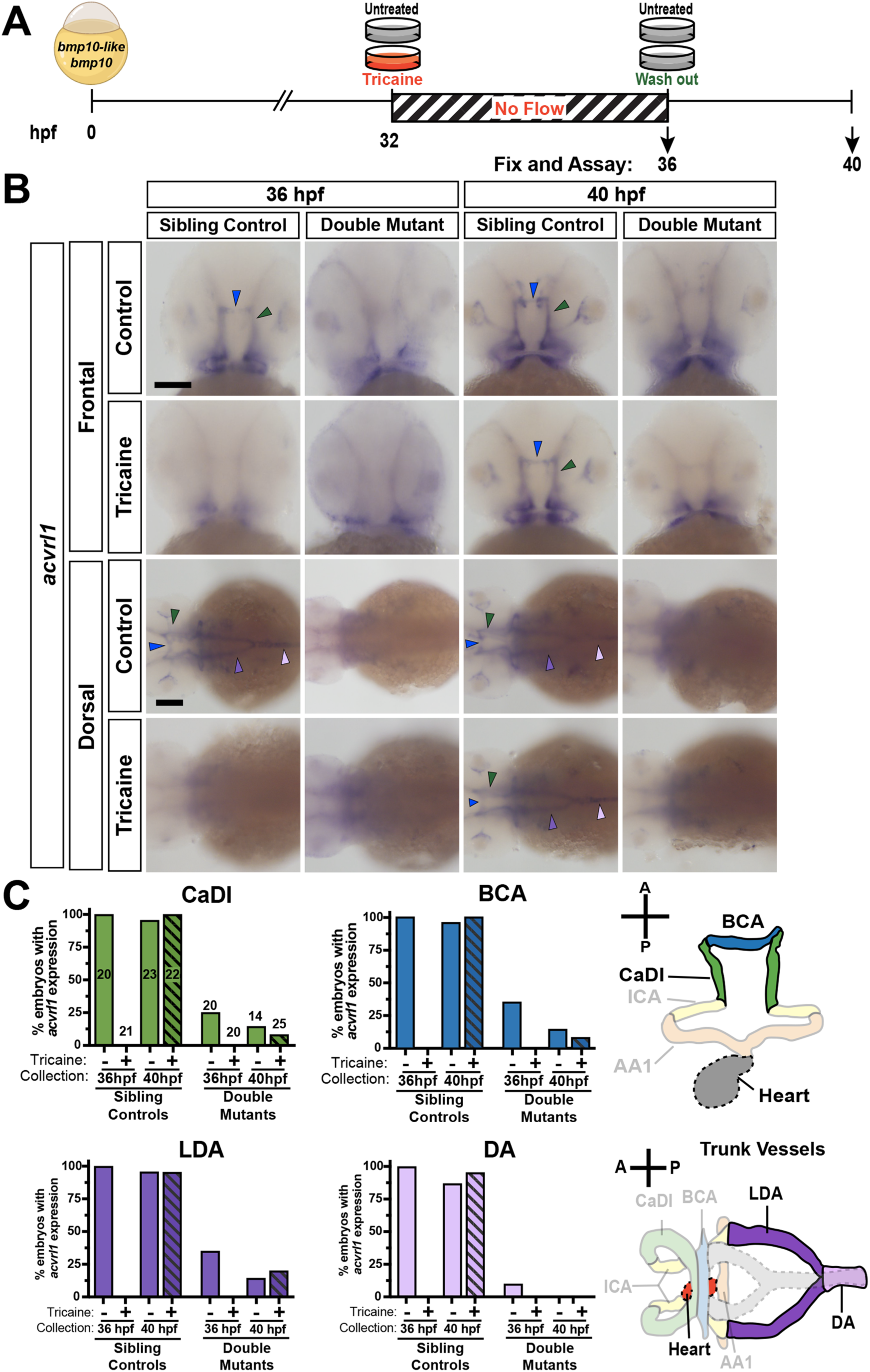
Bmp10 acts downstream of blood flow in maintaining *acvrl1* mRNA expression. **a.** Experimental design. At 32 hpf, embryos from a *bmp10^pt527/+^;bmp10-like^sa11654/sa11654^*incross were left untreated or treated with 800 µg/mL tricaine to stop blood flow, moved to fresh medium without tricaine at 36 hpf, and collected for *in situ* hybridization for *acvrl1* at 36 hpf (4 hours no flow) or 40 hpf (4 hours recovery). Non-tricaine treated embryos served as controls. **b**. Representative *acvrl1* images, frontal (top) and dorsal (bottom) views. Color-coded arrows mark presence of signal in indicated vessels. Scale bar = 100 µm. **c**. Percent of embryos with *acvrl1* expression detected in indicated vessels in sibling controls (*bmp10^+/+^;bmp10-like^-/-^*) and double mutants (*bmp10^-/-^;bmp10-like^-/-^*), without or with (hatched) tricaine treatment. The number of embryos analyzed per group, summed across 4 independent experiments, is displayed in CaDI graph. At right, schematics of vessels shown in frontal (top) and dorsal (bottom) views. Differences between groups for each vessel were computed by Fisher’s exact test and are reported in Supplementary Table 3. Color coding in **b,c:** caudal division of the ICA (CaDI, green), basal communicating artery (BCA, blue), lateral dorsal aorta (LDA, dark purple), dorsal aorta (DA, light purple).

### Bmp10/Alk1 activity is required for restoration of *acvrl1* expression in the absence of blood flow

To directly determine whether intravascular Bmp10 is sufficient to maintain *acvrl1* expression in the absence of blood flow, we used tricaine to stop heartbeat at 32 hpf in wild type embryos and immediately injected a local bolus of vehicle or recombinant human BMP10, which we previously demonstrated to activate Alk1 signaling in zebrafish [31], into the left CaDI. After 4 hours in a no-flow state (36 hpf), we assayed embryos by in situ hybridization (Fig. 4A). As expected, *acvrl1* was detected in all untreated embryos and was absent in all tricaine-treated, vehicle-injected embryos. By contrast, ∼75% of BMP10-injected embryos had detectable expression in and around the left CaDI (Fig. 4B). This percent rescue was similar to rescue of *cxcr4a* (∼66%), which is enhanced in the absence of flow and rescued to low basal levels by BMP10 injection (Fig. 4B and as previously reported [31]). No changes were observed in *cdh5* in any condition. These results suggest that Bmp10 is sufficient to maintain *acvrl1* expression in the absence of flow.

**Fig. 4.**
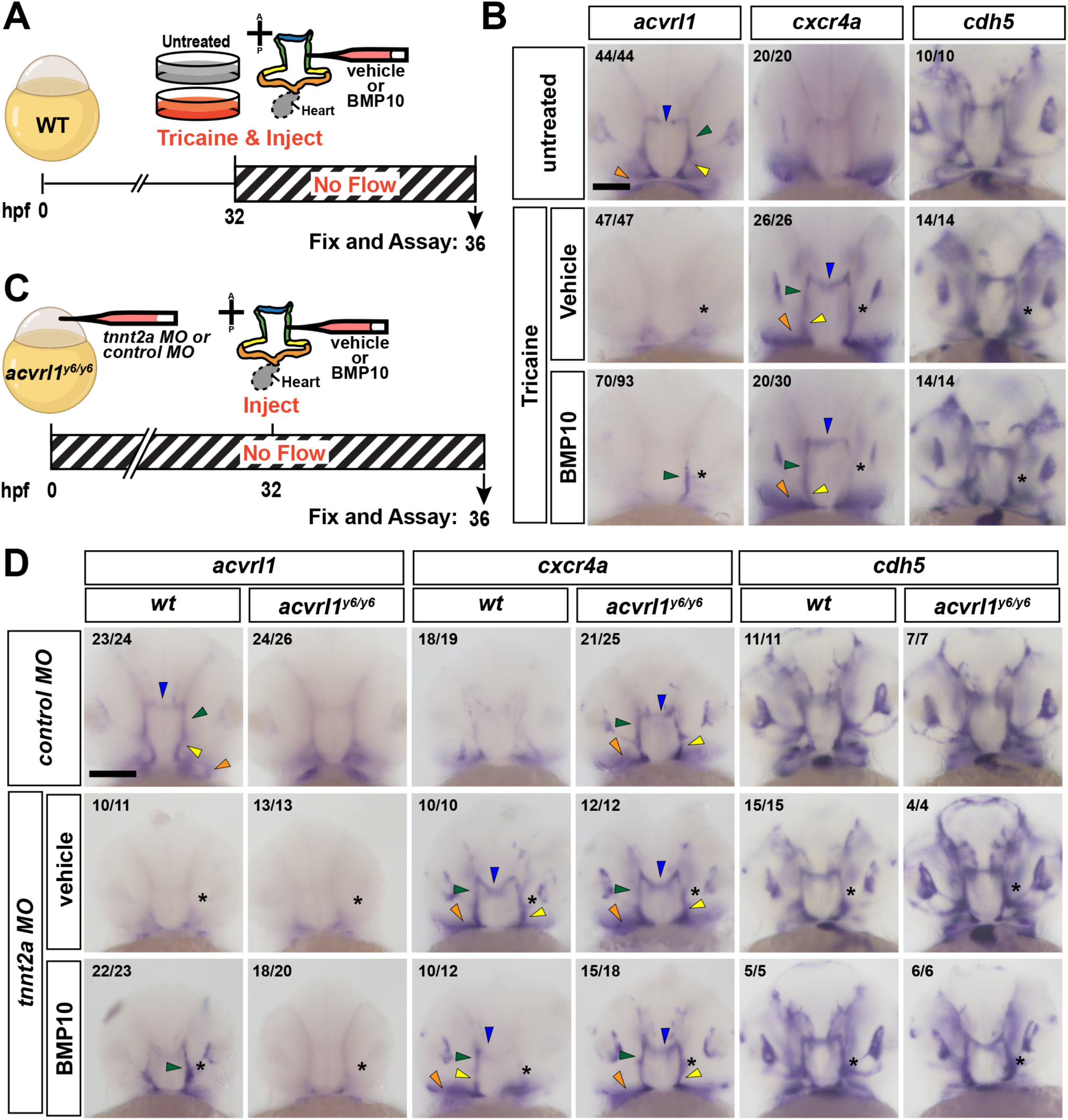
Bmp10/Alk1 activity is required for restoration of *acvrl1* expression in the absence of blood flow**. a.** Experimental design for panel b. At 32 hpf, wild type embryos were left untreated or treated with 1.6 mg/mL tricaine for 1-2 minutes to quickly stop heartbeat, transferred to 800 µg/mL tricaine, and immediately injected with vehicle or BMP10 in the left CaDI. Embryos were maintained in a no-flow state for 4 hours then fixed for assay at 36 hpf. **b**. Representative in situ hybridization signal for *acvrl1*, *cxcr4a*, and *cdh5* for indicated treatments. **c**. Experimental design for panel d. 1-cell stage embryos generated from an *acvrl1^+/y6^* incross were injected with control or *tnnt2a* morpholino at the 1-cell stage. *tnnt2a* morphants were injected with vehicle or BMP10 in the left CaDI at 32 hpf, and all embryos were fixed for assay at 36 hpf. **d.** Representative in situ hybridization for *acvrl1*, *cxcr4a*, and *cdh5* for indicated treatments and genotypes. In **b**,**d**: Asterisk indicates injection site; color-coded arrows indicate AA1 (orange), ICA (yellow), CaDI (green), and BCA (blue). Ratio in each panel, X/Y = number of embryos with represented phenotype/total assayed over 4 (**b**) or 5 (**d**) independent experiments. Frontal views, anterior at top. Scale bars = 100 µm

In a complementary approach, to directly assay the requirement for Alk1 activity in flow-mediated maintenance of *acvrl1* expression, we injected embryos derived from an *acvrl1^y6/+^* incross at the 1-cell stage with control or *tnnt2a* morpholino. At 32 hpf, we microinjected a local bolus of vehicle or recombinant human BMP10 into the left CaDI of *tnnt2a* morphants and fixed all embryos at 36 hpf for in situ hybridization (Fig. 4C). Embryos were scored blinded and genotyped after scoring; we present data from wild type siblings (*acvrl1^+/+^*) and *acvrl1^y6/y6^* mutants. *acvrl1* mutants injected with control morpholino (Fig. 4D, top row) showed decreased *acvrl1* expression, increased *cxcr4a* expression, and unchanged *cdh5* expression compared to wild type siblings. Similarly, *tnnt2a* morphants in which vehicle was injected into the CaDI at 32 hpf also showed decreased *acvrl1* expression, increased *cxcr4a* expression, and unchanged *cdh5* expression compared to wild type control morphants, regardless of *acvrl1* genotype (Fig. 4D, middle row). These data support our previous findings regarding the importance of Alk1 activity and blood flow in *acvrl1* expression [22]. Importantly, injection of BMP10 into the left CaDI of 32 hpf *tnnt2a* morphants locally increased *acvrl1* expression and decreased *cxcr4a* expression in wild type siblings, but not in *acvrl1* mutants, with no effect on *cdh5* expression (Fig. 4D, bottom row). Together, these data add support to the conclusion that Bmp10/Alk1 signaling is the key effector of “blood flow” that boosts *acvrl1* expression beyond basal levels, and suggest that in the absence of flow, low levels of residual Alk1 protein present at 32 hpf are sufficient to transduce the exogenous BMP10 signal. Moreover, because *acvrl1* is inducible in vessels that never experienced flow, we conclude that mechanical force is not required to prime this positive feedback regulatory mechanism.

### Establishment and validation of a transgenic zebrafish *acvrl1* reporter

Based on data demonstrating that mouse *Acvrl1* intron 2 (homologous to zebrafish *acvrl1* intron 1) contains critical regulatory elements that drive arterial gene expression [19], we set out to generate a zebrafish *acvrl1* reporter line. To this end, we injected into 1-cell stage embryos a plasmid containing a 1910-bp fragment of *acvrl1* (3 bases corresponding to the 3’ end of noncoding exon 1, plus 1907 bases corresponding to the 5’ end of intron 1) upstream of a basal promoter and *egfp*, and isolated a single stable transgenic line, *Tg(acvrl1e5:egfp)^pt517^* (Fig. 5A). Additionally, utilizing CRISPR/Cas9, we deleted a 1242 bp fragment from the 5’ end of the *Tg(acvrl1e5:egfp)^pt517^* transgene to generate a nested line, *Tg(acvrl1e5:egfp)^pt552^,* which retains 668 bp of *acvrl1* intron 1 (Fig. 5A). We mapped the *Tg(acvrl1e5:egfp)^pt517^* insertion to chromosome 20 within intron 6 of the 7-exon *pinx1* gene (ENSDARG00000023532), just upstream of *sox7* (Fig. 5A). Both *Tg(acvrl1e5:egfp)^pt517^* (Fig. 5B; Supplementary Fig. 5) and *Tg(acvrl1e5:egfp)^pt552^* (Fig. 5C) predominantly express EGFP in the arterial ECs that express endogenous *acvrl1*, including ECs in the aortic arches, ICA, CaDI, BCA, LDA, and DA, as well as the OA, PCS, and BA. Notably, expression is absent in *acvrl1*-negative arteries, such as the intersegmental and central arteries (Supplementary Fig. 5). Moreover, EGFP is absent in nearly all veins, although the remodeling caudal vein plexus expresses EGFP up until 2.5 dpf (Supplementary Fig. 5). These data strongly suggest that this *acvrl1* intron 1 segment directs gene expression in a manner similar to endogenous *acvrl1* control elements. However, we cannot rule out the possibility that this transgene co-opted local regulatory elements that are permissive for endothelial gene expression: *sox7* (but not *pinx1*) is expressed in arterial ECs, albeit in a broader pattern that includes the intersegmental arteries (not shown), and it is also expressed in the hindbrain (Fig. 5C). Accordingly, we conclude that this *acvrl1* intron 1 fragment can drive expression in or limit gene expression to arterial ECs, establishing a gene expression pattern that closely resembles endogenous *acvrl1*.

**Fig. 5.**
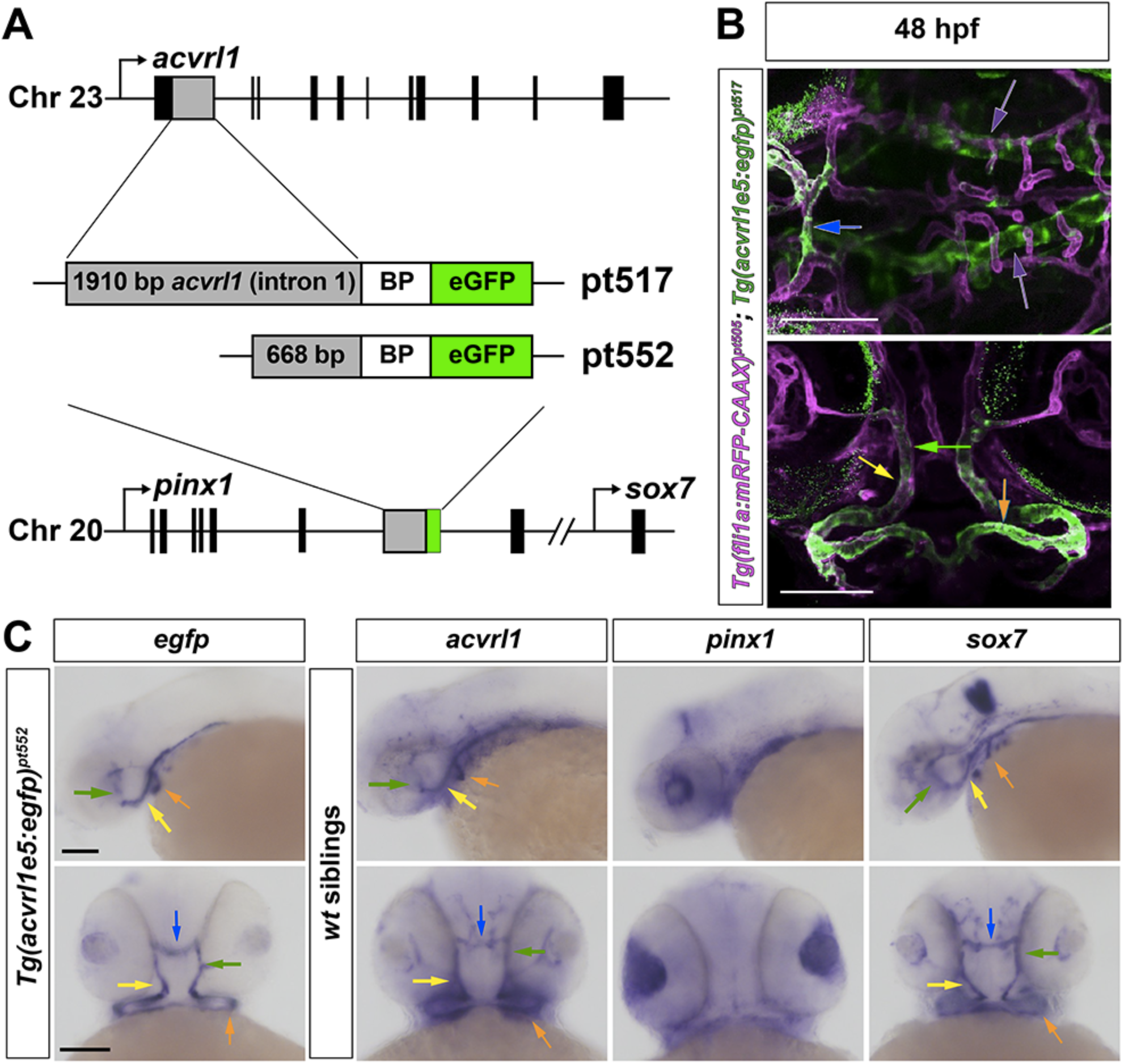
Establishment and validation of transgenic zebrafish *acvrl1* reporter. **a** Schematics showing transgene configurations and genomic insertion site of *Tg(acvrl1e5:egfp)^pt517^*and *Tg(acvrl1e5:egfp)^pt552^*. **b** 2D projections of two-photon/confocal z-series of *Tg(acvrl1e5:egfp)^pt517^* (green) and *Tg(fli1a:mRFP-CAAX)^pt505^* (magenta) at 48 hpf. Top panel: dorsal view, anterior left. Bottom panel: frontal view, anterior top. Scale bar = 100 µm. See Supplementary Fig. 5 for extensive characterization of *Tg(acvrl1e5:egfp)^pt517^*, 1.5-5 dpf. **c** In situ hybridization signal for *egfp*, *acvrl1*, *pinx1*, and *sox7*, in *Tg(acvrl1e5:egfp)^pt552^* (*egfp*) or non-transgenic siblings at 48 hpf. Colored arrows indicate AA1 (orange), ICA (yellow), CaDI (green), and BCA (blue). Top: lateral view, anterior left. Bottom: frontal view, anterior top. Scale bars = 100 µm

### Blood flow regulates *acvrl1* at the level of transcription

To determine whether regulation of *acvrl1* mRNA expression by flow-mediated Bmp10/Alk1 signaling occurs at the transcriptional level, we took advantage of our *Tg(acvrl1e5:egfp)^pt552^* reporter. We used tricaine to stop heartbeat at 44 hpf in *Tg(acvrl1e5:egfp)^pt552^*embryos and immediately injected a local bolus of vehicle or recombinant human BMP10 into the left CaDI. After 4 hours in a no-flow state (48 hpf), we assayed embryos by in situ hybridization (Fig. 6A). Expression of *egfp* in AA1 and ICA was refractory to loss of blood flow with vehicle injection, whereas *egfp* expression in CaDI and BCA was lost, similar to effects on endogenous *acvrl1* (Fig. 6B). Expression of *egfp* in *Tg(acvrl1e5:egfp)^pt517^*behaved similarly in response to stopped flow (A. Biery Kunz, data not shown). In accordance with results for endogenous *acvrl1*, in ∼ 78% of embryos, microinjection of BMP10 maintained *egfp* expression in the left CaDI in the absence of flow (Fig. 6B). No changes were observed in *cdh5* in any condition (Fig. 6B).

**Fig. 6.**
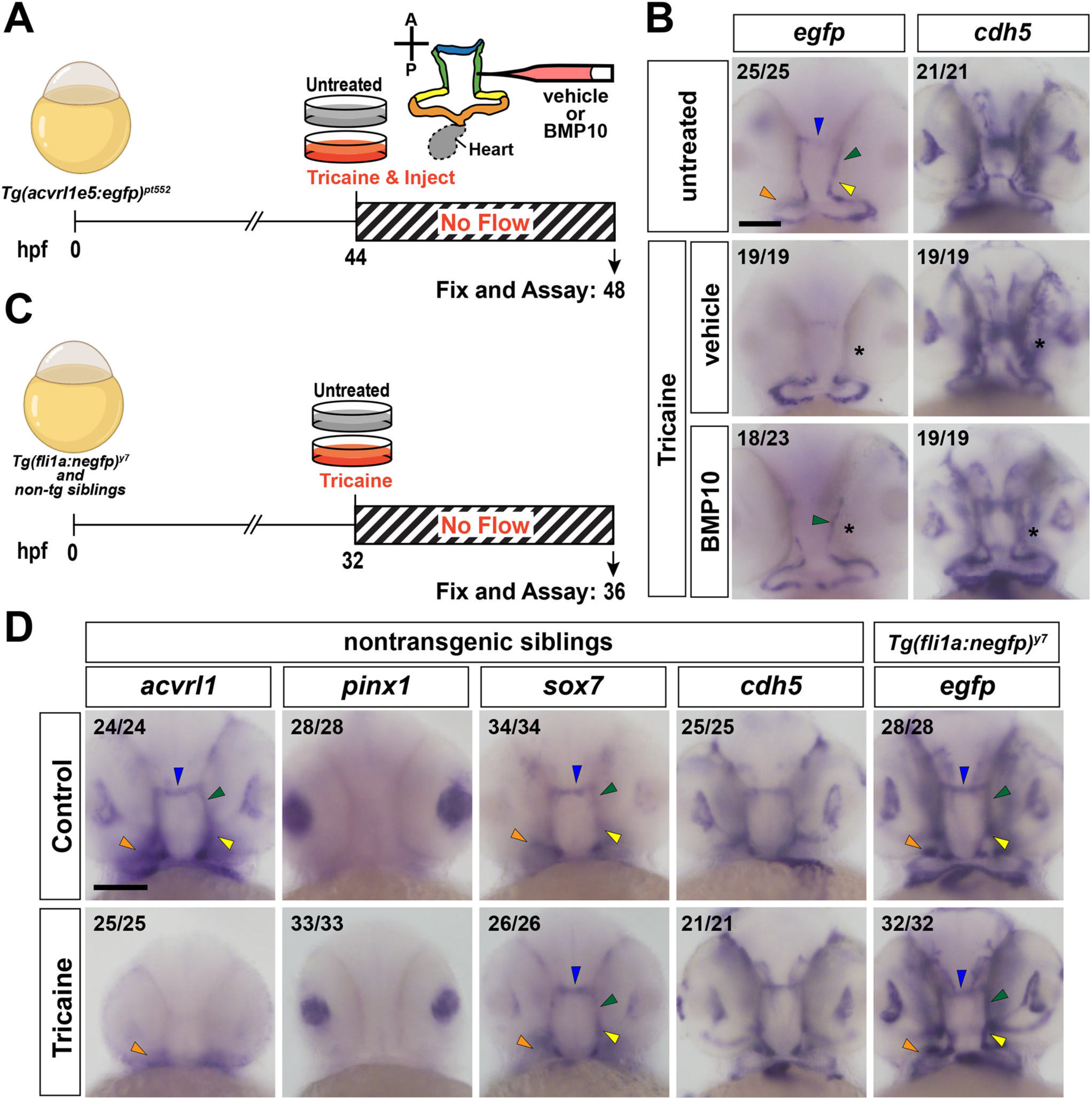
Blood flow regulates *acvrl1* expression at the level of transcription. **a** Experimental design for panel **b**. At 44 hpf, *Tg(acvrl1e5:egfp)^pt552^* embryos were left untreated or treated with 1.6 mg/mL tricaine for 1-2 minutes to quickly stop heartbeat, transferred to 800 µg/mL tricaine, and immediately injected with vehicle or BMP10 in the left CaDI. Embryos were maintained in a no-flow state for 4 hours then fixed for assay at 48 hpf. **b** Representative in situ hybridization for *egfp* and *cdh5* for indicated treatments. Asterisk indicates injection site. Ratio in each panel, X/Y = number of embryos with represented phenotype/total assayed over 3 independent experiments. **c** Experimental design for panel **d**. At 32 hpf, *Tg(fli1a:negfp)^y7^* or non-transgenic siblings were left untreated or treated with 800 µg/mL tricaine to stop blood flow and collected at 36 hpf (4 hours no flow). **d** Representative in situ hybridization for *acvrl1, pinx1*, *sox7*, *cdh5,* and *egfp*. Ratio in each panel, X/Y = number of embryos with represented phenotype/total assayed over 2 independent experiments. Color coding in **a**,**b,d**: AA1 (orange), ICA (yellow), CaDI (green), and BCA (blue). Frontal views, anterior top. Scale bars = 100 µm

To ensure that the loss of *egfp* expression was not reflective of flow sensitivity of *pinx1* or *sox7*, which flank the *acvrl1e5:egfp* transgene insertion site on chromosome 20, nor of instability of *egfp* transcript, we assayed embryos derived from an unrelated hemizygous EC transgenic, *Tg(fli1a:negfp)^y7^*, outcrossed to wild type. We treated transgenic and non-transgenic embryos with 800 μg/mL tricaine at 32 hpf and assayed gene expression at 36 hpf via in situ hybridization (Fig. 6C). Stopped flow decreased *acvrl1* expression, as expected, but had no effect on *pinx1, sox7, cdh5* (assayed in non-transgenic siblings) or *egfp* expression (Fig. 6D). Together, these data suggest that Bmp10/Alk1 activity regulates *acvrl1* transcription at least in part via a *cis* element in *acvrl1* intron 1.

### Shear stress induces *ACVRL1* in cultured human endothelial cells in an ALK1 ligand-dependent manner

Collectively, our zebrafish data demonstrate that Bmp10 acts downstream of blood flow to regulate *acvrl1* expression at the level of transcription. To determine whether this relationship holds true in human ECs, we first assayed *ACVRL1* expression in human umbilical vein ECs (HUVECs) cultured in complete (2% serum-containing) medium and subjected to a range of shear stress magnitudes (2, 8, 14, and 20 dyn/cm^2^) for 28 hours (Fig. 7A). Compared to static conditions, *ACVRL1* was significantly increased at all shear stress magnitudes, with highest induction at 8 dyn/cm^2^. By contrast, the flow-sensitive gene, *KLF2*, displayed a clear magnitude-dependent increase from 2-20 dyn/cm^2^ (Fig. 7A). These data suggest that *ACVRL1* expression is maximally induced by moderate shear stress magnitudes via a mechanism that is likely different from *KLF2*.

**Fig. 7.**
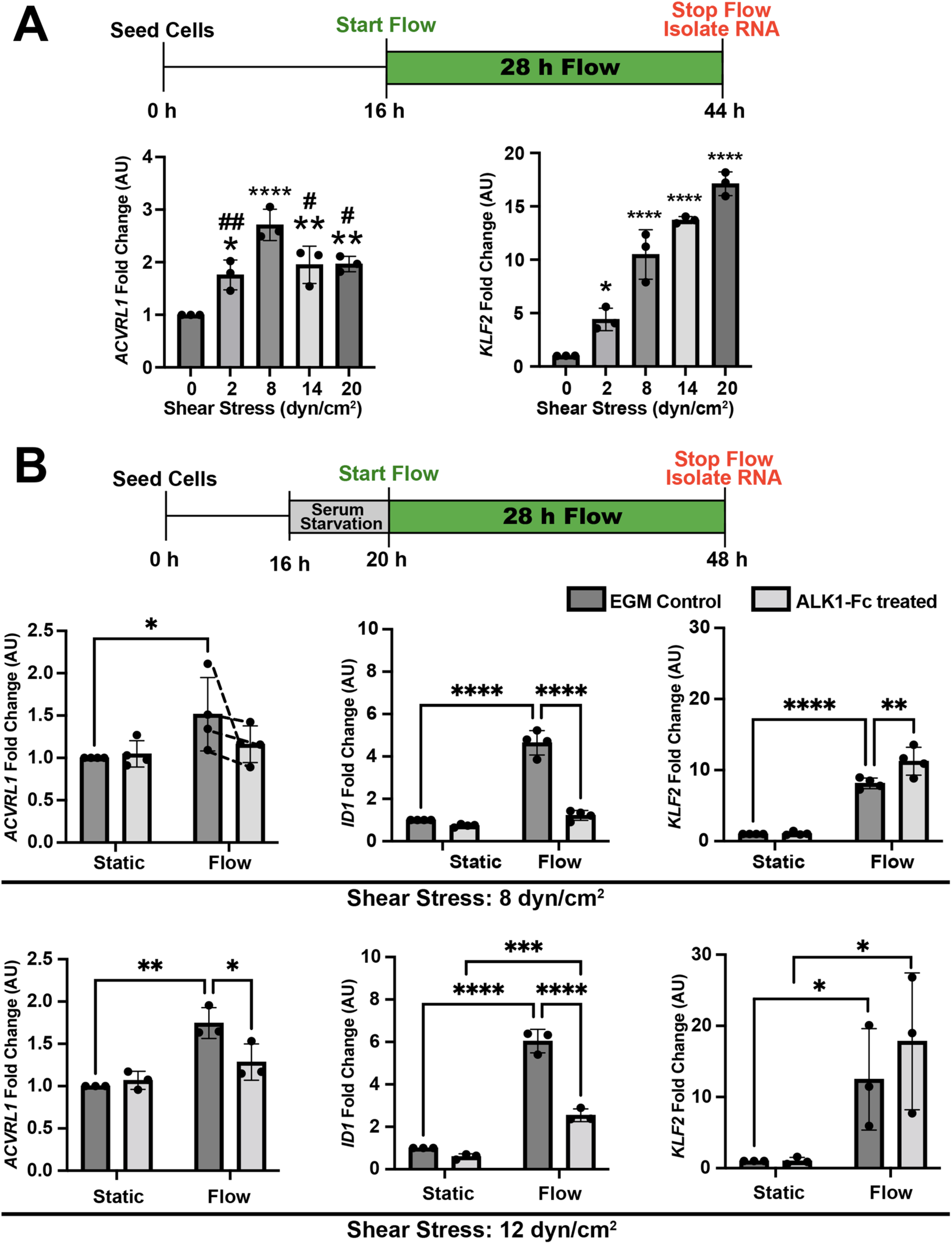
Shear stress-induced *ACVRL1* upregulation in human endothelial cells is dependent on ALK1 ligands. **a** HUVECs were seeded into microfluidic devices for a 16-hour acclimation period and subjected to varying magnitudes of shear stress for 28 hrs. RNA was isolated and RT-qPCR was performed for indicated genes. n = 3 independent experiments. Differences between groups were computed by one-way ANOVA with Tukey’s post-hoc analysis for comparisons between all groups. Comparisons made to static control: *p<0.05, **p<0.01, ***p<0.001, ****p<0.0001. Comparisons made to 8 dyn/cm^2^: # p<0.05, ##p<0.01. **b** HUVECs were seeded into microfluidic devices for a 16-hour acclimation period followed by a 4-hour serum starvation period. Shear stress, 8 or 12 dyn/cm^2^, was applied for 28 hours in media containing FBS pre-treated with ALK1-Fc. RNA was isolated and RT-qPCR was performed for indicated genes. n = 3 (12 dyn/cm^2^) or 4 (8 dyn/cm^2^) independent experiments. Two-way ANOVA was used to detect differences across flow conditions and across media treatments. Multiple comparisons between flow conditions and media treatments were computed using Tukey’s post-hoc analysis. *p< 0.05, **p<0.01, ***p<0.001, ****p<0.0001

To determine whether flow-induced upregulation of *ACVRL1* is dependent on ALK1 ligands in FBS, we assayed HUVEC response to flow in medium containing ALK1-Fc-treated (and therefore ALK1 ligand -depleted) serum [38]. Use of ligand-depleted serum abrogated *ACVRL1* induction in response to 8 or 12 dyn/cm^2^ shear stress at 28 hours but had no effect on basal *ACVRL1* expression in static cultures (Fig. 7B). Similarly, shear stress-mediated induction of a known downstream target of ALK1 signaling, *ID1,* was significantly decreased under ligand-depleted conditions. By contrast, *KLF2* was induced in the presence or absence of ALK1 ligands, with slightly higher expression under ligand-depleted conditions (Fig. 7B). Together, these data suggest that *ACVRL1* is modestly increased by shear stress in HUVECs in an ALK1 ligand-dependent manner, supporting our zebrafish data, and that ALK1 signaling may dampen shear stress-induced *KLF2*.

## Discussion

Blood flow exerts mechanical forces on the vessel wall and exposes ECs to a myriad of circulating factors. Understanding how interactions between physical forces and signaling drive EC gene expression and behavior is critical for understanding the pathogenic mechanisms of HHT and other diseases. Using a zebrafish model, we previously reported that *acvrl1* expression depends on blood flow [22]. In this work, we deepen our mechanistic understanding of this phenomenon. We demonstrate that *acvrl1* expression initiates but fails to be maintained in the complete absence of blood flow, with some arterial ECs (within the CaDI, BCA, LDA and DA) more sensitive to the lack of flow than other arterial ECs (AA1, ICA). Additionally, using *bmp10;bmp10-like* double mutants, we demonstrate a similar dependence of *acvrl1* expression on the Alk1 ligand, Bmp10. We also demonstrate that *acvrl1* expression is rapidly downregulated in the absence of flow and rapidly restored when flow resumes; moreover, Bmp10/Alk1 signaling is necessary for *acvrl1* restoration with resumed flow and sufficient for *acvrl1* induction in the absence of flow. Finally, using a zebrafish *acvrl1:egfp* reporter line, we demonstrate that flow- and Bmp10/Alk1-mediated regulation of *acvrl1* expression occurs at the level of transcription. Thus, we conclude that *acvrl1* expression is regulated by a positive feedback mechanism that requires Bmp10/Alk1 activity.

In zebrafish, *acvrl1* is expressed most prominently in AA1 and the ICA, which are the cranial arteries closest to the heart, and *acvrl1* expression within these ECs is relatively refractory to loss of blood flow. Given that Bmp10/Alk1 signaling induces *acvrl1* expression and that *bmp10* and *bmp10-like* are expressed in the embryonic heart [31], a gradient of circulating Bmp10, with local diffusion in the absence of flow, might explain the high expression in and reduced flow sensitivity of AA1 and ICA ECs. However, the fact that *acvrl1* expression in these ECs is similarly refractive to the loss of flow or Bmp10, and the observation that there is no BMP10 gradient in human plasma [39], fail to support this idea. Accordingly, we hypothesize that basal levels of *acvrl1* are maintained in a flow- and Bmp10-independent manner by an unknown mechanism that may be most robust in AA1 and ICA ECs. Notably, the flow- and Bmp10-refractory AA1 and proximal ICA originate from bilateral angioblast clusters known as midbrain organizing centers, whereas the flow- and Bmp10-sensitive CaDI and BCA originate from rostral organizing centers [40]. As heterogeneity exists within arterial EC populations [41], it is possible that *acvrl1* expression is regulated somewhat differently in ECs derived from different embryonic lineages. How lineage-specific genetic programs might result in differentially regulated *acvrl1* expression requires further investigation.

Our results strongly suggest that blood flow distributes ligands to Alk1 to induce a positive feedback loop in which Alk1 signaling enhances its own transcription. In zebrafish embryos, visibly decreased *acvrl1* expression is discernible by 1-2 hours after stopping blood flow. As the average half-life of maternally expressed mRNA in zebrafish is ∼1-4 hours [42], this time scale of loss reflects expected rates of decay after transcriptional downregulation. While decay rates are influenced by codon usage, 3’ UTR sequence, poly(A) tail length, or active miRNA-mediated destabilization [42–44], flow stoppage resulted in similar kinetics of *egfp* transcript loss in our *acvrl1:egfp* reporter line, suggesting that sequence-specific determinants of decay rate play a minimal role in *acvrl1* transcript loss.

A balance between positive and negative feedback mechanisms is required for controlling TGF-β family signaling pathways [45] and there is precedence for positive feedback in BMP signaling during development [46]. With respect to *ACVRL1* expression, this mechanism has received little attention to date, but is supported by some published work. For example, treatment of human umbilical artery ECs with BMP9 (5 ng/mL for 48 hours) induced *ACVRL1* expression [47], and BMP4/ALK2 signaling through phosphorylated SMAD1, which also relays signal from BMP10/ALK1, induces *ACVRL1* in human aortic ECs [48].

How flow- and ligand-mediated Alk1 activation regulates *acvrl1* transcription is unknown. To begin to address this question, we employed a transgenic approach to identify flow-controlled regulatory elements in zebrafish *acvrl1* intron 1. We chose this region because mouse *acvrl1* intron 2, which is homologous to zebrafish and human intron 1, is required to drive arterial-specific transgene expression [18,19]. Additionally, compared to downstream introns, first introns are generally more conserved across species and harbor a relative abundance of *cis* regulatory elements and active chromatin marks [49]. Indeed, ENCODE data (accessed via the UCSC Genome Browser, http://genome.ucsc.edu, GRCh38/hg38) reveal a broad H3K27Ac peak across human *ACVRL1* intron 1 [50,51]. Using a plasmid in which the 5’ half of zebrafish *acvrl1* intron 1 drives *egfp* expression from a basal heterologous promoter, we successfully isolated a single transgenic zebrafish line that recapitulated the flow responsiveness of endogenous *acvrl1.* However, additional independent stable lines exhibited non-EC transgene expression, suggesting strong positional effects. The transgene in *Tg(acvrl1e5:egfp)^pt517^* and the subline, *Tg(acvrl1e5:egfp)* ^pt552^, inserted into the final intron of *pinx1* exon 6, which is just upstream of the arterial EC-expressed gene, *sox7*. Based on HUVEC ENCODE data showing six correlated H3K27Ac/GATA2 ChIP seq peaks in *PINX1* intron 6 and conserved synteny in this region between human (chromosome 8) and zebrafish (chromosome 20), we surmise that this transgene coopted regulatory elements that allow EC-specific expression. Supporting this conclusion, additional transient transgenic expression studies using the smaller *pt552 acvrl1* intron 1 element in conjunction with the zebrafish *acvrl1* promoter failed to drive EC transgene expression (data not shown). Accordingly, we conclude that this *acvrl1* intron 1 element is sufficient to confer transgene flow responsiveness, but it is not sufficient to direct EC-specific expression. While canonical Alk1 signaling acts via phosphorylated Smad1, Smad5, or Smad9, the *cis* element driving *egfp* expression in *Tg(acvrl1e5:egfp)* ^pt552^ has no TRANSFAC-predicted Smad1/5/9 binding sites [52]. Current efforts are focused on assaying necessity of *acvrl1* intron 1 elements via CRISPR/Cas9-mediated deletion and defining trans-acting factors that function in critical regions.

Although we have demonstrated that BMP10 can serve as a proxy for blood flow in induction of *acvrl1* expression, it remains possible that blood flow/shear stress might modulate *acvrl1* expression via additional mechanisms. Blood flow dramatically alters chromatin state in ECs, inducing *KLF2* and *KLF4* expression [53,54] and increasing accessibility of KLF family transcription factor binding sites [55], and KLF2, KLF4, and KLF6 are predicted to strongly bind to the human *ACVRL1* promoter (MATCH Suite, TRANSFAC 2.0 release 2.3). However, KLF2/4 activity is likely not required for BMP10-induced *acvrl1* expression in zebrafish, as we demonstrated that intravascularly injected BMP10 induces endogenous *acvrl1* in embryos that never experience blood flow (*tnnt2a* morphants). Moreover, we previously demonstrated that *klf2a* knockdown has no effect on *acvrl1* expression [22]. By contrast, it remains possible that vascular injury-induced expression of a *KLF6* paralog [16] might prime BMP10 induction of *acvrl1* in these embryos.

Our results in HUVECs partially corroborate our findings in zebrafish embryos, but important differences warrant discussion. In HUVECs, we demonstrated that shear stress induced modest increases in *ACVRL1* expression with no clear relationship between shear stress magnitude (2-20 dyn/cm^2^) and response, supporting our previous findings that lowered viscosity (and thus decreased shear stress) did not affect *acvrl1* expression in zebrafish embryos [22]. Moreover, in HUVECs, shear stress-induced *ACVRL1* was abrogated by sequestration of serum-borne ALK1 ligand using ALK1-Fc. These data support the idea that ALK1 ligand serves as a proxy for blood flow in this response. Notably, however, in cultured ECs, shear stress was required for priming a response to ALK1 ligand. That is, we were not able to induce *ACVRL1* expression in cultured ECs (HUVECs, human pulmonary artery ECs, or human dermal microvascular ECs) by stimulation with 2% serum, BMP9, or BMP10 (0.06 – 7.5 ng/mL) for various times (0.5 to 24 hours) under static conditions (data not shown). It is possible that in culture, shear stress is required to alter chromatin structure to allow ligand-mediated *acvrl1* induction and/or that variables such as substrate stiffness or extracellular matrix composition prevent full recapitulation of an in vivo response.

Further work is required to understand the relationship between KLF2 and ALK1. In contrast to *ACVRL1*, *KLF2* expression exhibited a robust positive correlation with shear stress magnitude (2-20 dyn/cm^2^) that was not dampened by sequestration of ALK1 ligands. In fact, we consistently saw a modest increase in *KLF2* expression in sheared HUVECs in the absence versus presence of ALK1 ligands. This finding suggests that ALK1 signaling might actually dampen *KLF2* response to shear stress, perhaps serving as a negative feedback mechanism to constrain *KLF2*.

The positive feedback mechanism that governs *ACVRL1* expression can be exploited for treatment of patients with vascular malformations. Because *Acvrl1* overexpression rescues AVM development in *Eng* null mice [14], biologics or small molecules that enhance ALK1 activity and increase *ACVRL1* expression may be beneficial in HHT1 patients, in whom *ACVRL1* is intact. Moreover, if ALK1 signaling does in fact dampen *KLF2* expression, these therapies might also be useful in treatment of cerebral cavernous malformations, which are driven in part by increased expression of *KLF2* [56].

## Acknowledgements

We thank U. Sonmez (Carnegie Mellon University, Pittsburgh, PA, USA) for mesofluidic device design and mold fabrication; Z. Kupchinsky, D. Wright, and J. Fiore (University of Pittsburgh, Pittsburgh, PA, USA) for excellent fish care; E. Rochon, M. Schubert, J.S. Yang (University of Pittsburgh) for technical contributions; Koichi Kawakami (National Institute of Genetics, Mishima, Japan) for permission to use Tol2-mediated transgenesis in zebrafish; A. Hinck (University of Pittsburgh) for generating recombinant human BMP10; K. Przyklenk and R. Thomas (Central Michigan University, Mount Pleasant, MI, USA) for guidance on statistical analysis of Bmp10 and flow interactions in zebrafish embryos.

## Author contributions

Anthony Anzell, Amy Biery Kunz, and Beth Roman conceptualized and designed the studies. Sarah Young, Thanhlong Tran, and Xinyan Lu generated the *acvrl1-*reporter transgenic zebrafish lines. Anthony Anzell, Amy Biery Kunz, James Donovan, and Beth Roman contributed to the acquisition, analysis, and interpretation of the data. Anthony Anzell and Beth Roman wrote the original draft. All authors reviewed and approved the final manuscript for publication.

## Compliance with ethical standards

All procedures performed in studies involving animals were in accordance with ethical standards of the University of Pittsburgh (PHS approval A3187-01)

