## Supplemental Material for "Blood flow regulates *acvrl1* transcription via ligand-dependent Alk1 activity"

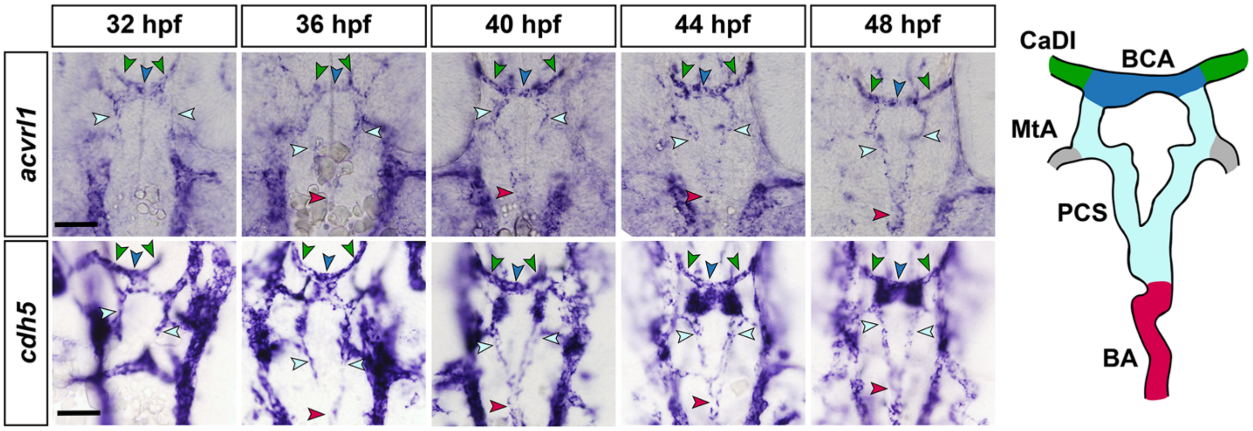


**Supplementary Fig. 1** *acvrl1* expression in the central cranial arterial vasculature. In situ hybridization for *acvrl1* (top, using a sensitive 3-riboprobe combination) or *cdh5* (bottom) in wild type embryos, 32-48 hpf. Vibratome sections (*acvrl1* = 50 µm; *cdh5* = 100 µm), dorsal view, anterior up. Color coding (arrows and vessel schematic): caudal division of the internal carotid artery (CaDI, green), basal communicating artery (BCA, dark blue), posterior communicating segments (PCS, light blue), basilar artery (BA, red). The BCA and PCS comprise the circle of Willis in zebrafish. MtA = metencephalic artery. Scale bars = 100 µm. Images representative of 3-8 embryos per probe per time point.

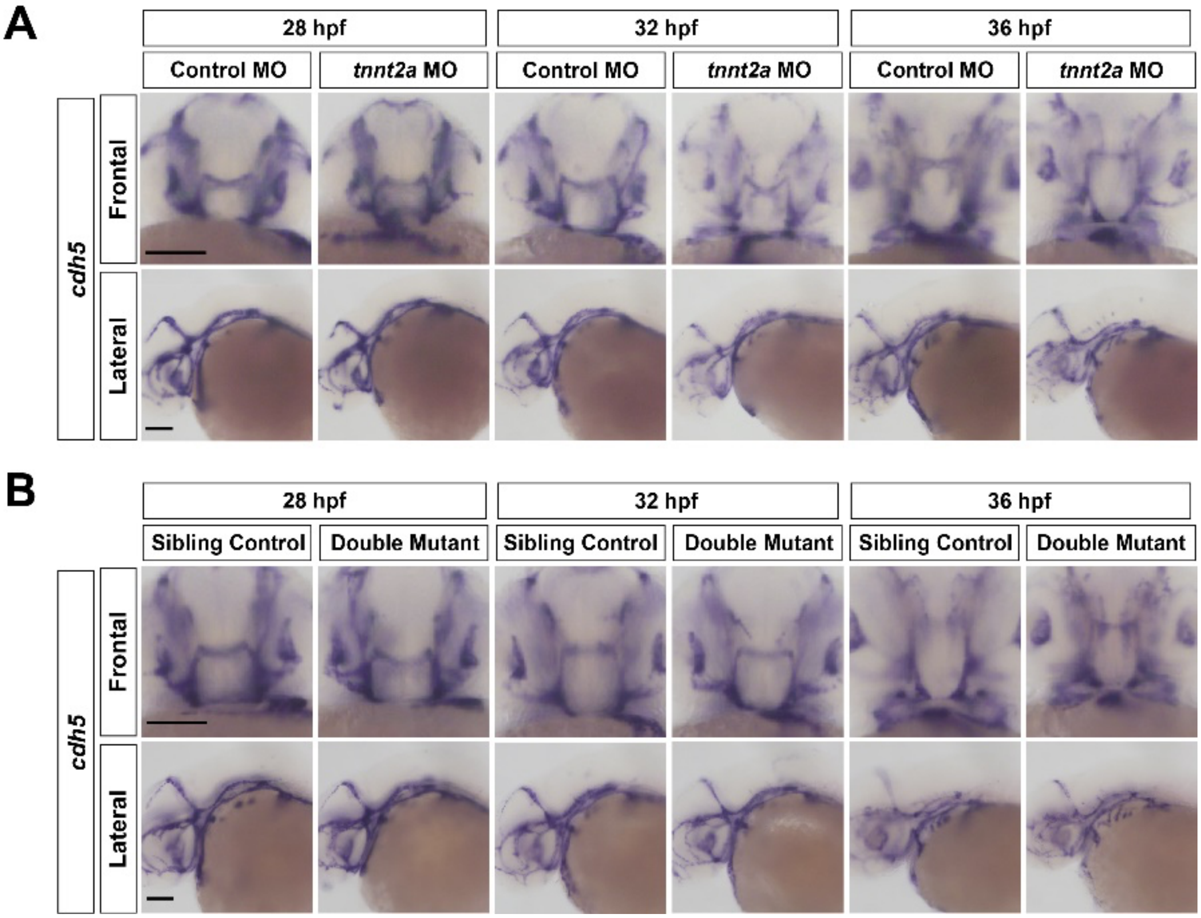


**Supplementary Fig. 2** *cdh5* expression is unaffected by loss of blood flow or *bmp10*. **a** Controls related to Fig.1. Wild type embryos were injected at the 1-cell stage with *tnnt2a* or control morpholino, fixed at indicated timepoints, and assayed by in situ hybridization for *cdh5*. n ≥ 21 embryos/group over 3 independent experiments. **b** Controls related to Fig. 2. Embryos generated from a $bmp10^{pt527/+}$;$bmp10\text{-}like^{sa11654/sa11654}$ incross were fixed at the indicated timepoints and assayed by in situ hybridization for *cdh5*. n ≥ 8 embryos/group over 3 independent experiments. Scale bars = 100 μm.

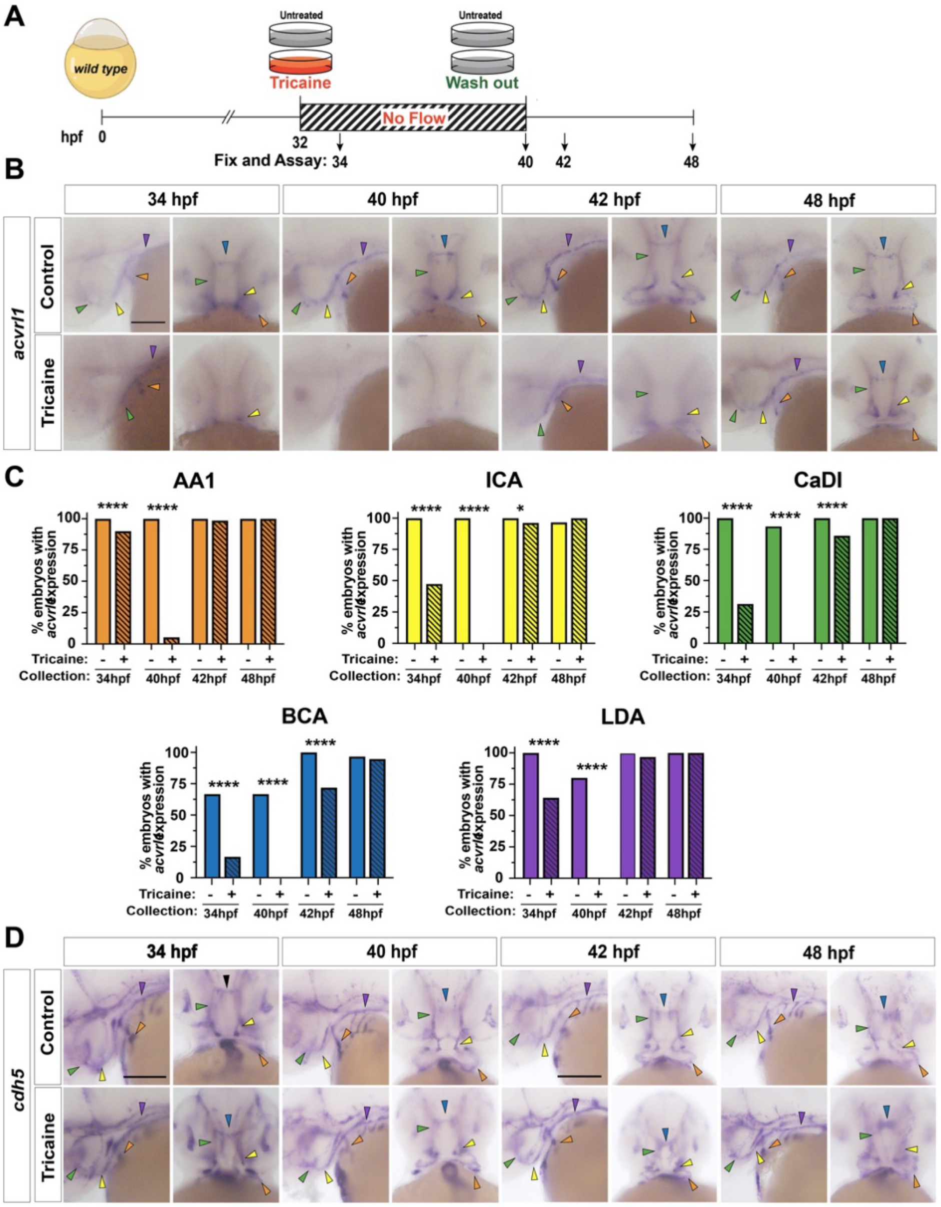


**Supplementary Fig. 3** *acvrl1* expression is sensitive to cessation and reinitiation of blood flow. **a** Experimental design. At 32 hpf, wild type embryos were treated with 800 μg/mL tricaine to stop blood flow. Tricaine was washed out at 40 hpf and embryos were allowed to recover for up to 8 hrs. Embryos were fixed at indicated time points and assayed by in situ hybridization. **b** Representative *acvrl1* images, lateral (first column) and frontal (second column) views. Color-coded arrows mark presence of signal in indicated vessels. Scale bar = 100 μm. **c** Percent of embryos with *acvrl1* expression detected in indicated vessels in control and tricaine treated embryos. n ≥ 50 embryos per group over 3 independent experiments. Differences between groups within each vessel were computed by Fisher's exact test *p<0.05, ****p<0.0001. **d** Representative *cdh5* images, lateral (first column) and frontal (second column) views. Color-coded arrows mark presence of signal in indicated vessels. Scale bar = 100 μm. Color coding: first aortic arch (AA1, orange), internal carotid artery (ICA, yellow), caudal division of the ICA (CaDI, green), basal communicating artery (BCA, blue), lateral dorsal aorta (LDA, dark purple).

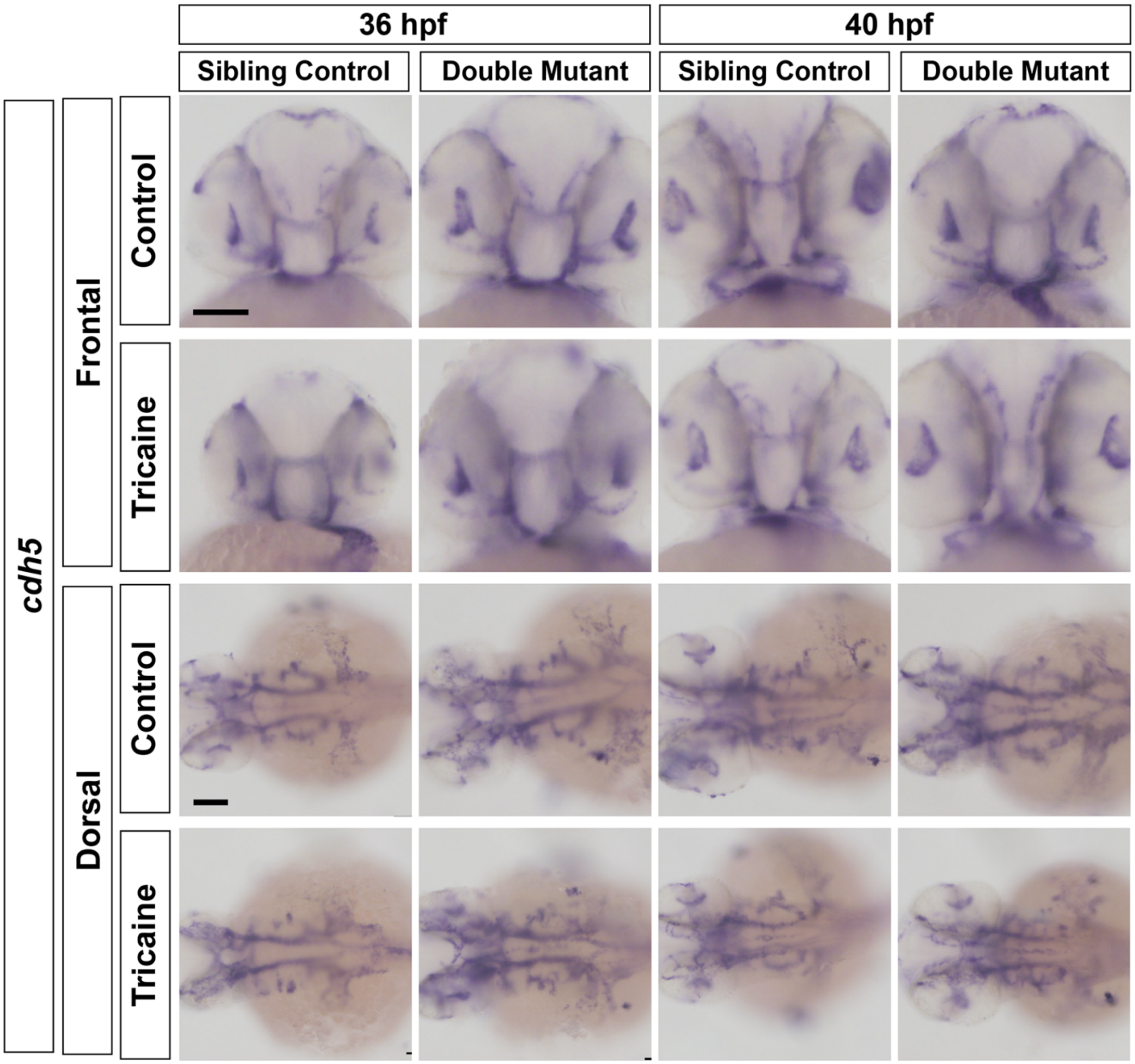


**Supplementary Fig. 4** Loss of blood flow or Bmp10 has no effect on *cdh5* mRNA expression. Controls related to Fig. 3. At 32 hpf, embryos from a $bmp10^{pt527/+}$/$bmp10\text{-}like^{sa11654/sa11654}$ incross were left untreated or treated with 800 μg/mL tricaine to stop blood flow, moved to fresh medium without tricaine at 36 hpf, and collected for in situ hybridization for *cdh5* at 36 hpf (4 hours no flow) or 40 hpf (4 hours recovery). Non-tricaine treated embryos served as controls. *cdh5* images, frontal (top) and dorsal (bottom) views of representative sibling controls ($bmp10^{+/+}$;$bmp10\text{-}like^{-/-}$) and double mutants ($bmp10^{-/-}$;$bmp10\text{-}like^{-/-}$). n ≥ 14 embryos/group across 4 independent experiments. Scale bars = 100 μm.

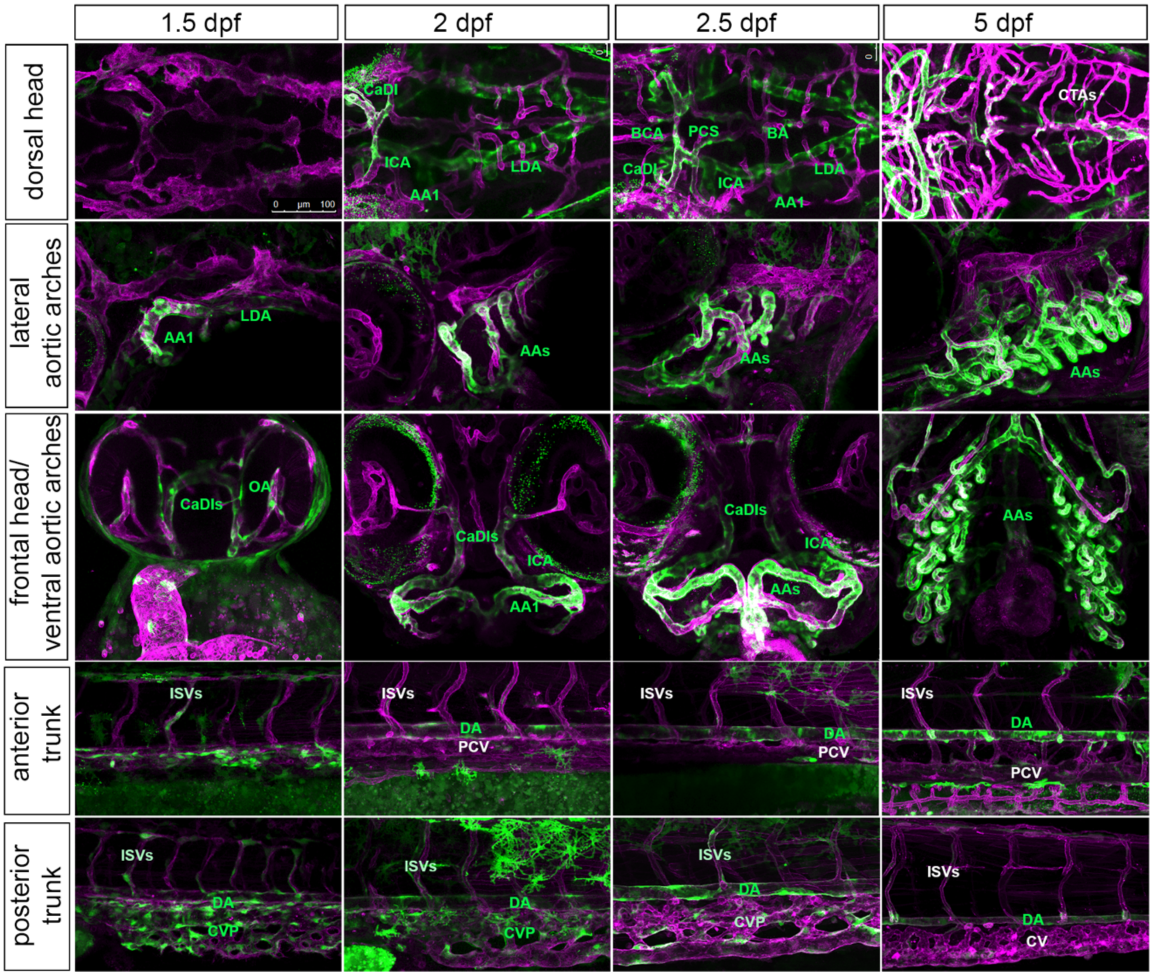


**Supplementary Fig. 5** EGFP expression in *acvrl1* reporter embryos faithfully recapitulates *acvrl1* expression. Double transgenic *Tg*(*acvrl1e5:egfp*)$^{pt517}$;*Tg*(*fli1a:mRFP*)$^{pt505}$ embryos and larvae, 1.5-5 days post-fertilization (dpf). Green, *acvrl1* EGFP reporter; magenta, pan-endothelial mRFP. Arterial vessels expressing EGFP are annotated in green font: AA1, first aortic arch; AAs, aortic arch arteries; BA, basilar artery; BCA, basal communicating artery; CaDI, cadual division of the internal carotid artery; DA, dorsal aorta; ICA, internal carotid artery; LDA, lateral dorsal aorta; OA, opercular artery; PCS, posterior communicating segments. Intersegmental vessels (ISV), which sprout from the *acvrl1*-positive DA, are weakly EGFP-positive after sprouting but lose expression over time. Central arteries (CTA), which sprout from *acvrl1*-negative veins, are transgene-negative. Veins are transgene-negative apart from the caudal vein plexus (CVP); expression is lost in this vessel as remodeling progresses to generate the caudal vein (CV). Scale bar = 100 µm.

| Time of Collection | Embryo | Vessel Segment | | | | | |
|---|---|---|---|---|---|---|---|
| | | AA1 | ICA | CaDI | BCA | LDA | DA |
| 28 vs 32 hpf | *control* MO | ns | ns | ns | ns | ns | ns |
| 28 vs 36 hpf | *control* MO | ns | ns | ns | ns | ns | ns |
| 32 vs 36 hpf | *control* MO | ns | ns | ns | ns | ns | ns |
| 28 vs 32 hpf | *tnnt2a* MO | ns | ns | ns | ns | p<0.001 | p=0.0054 |
| 28 vs 36 hpf | *tnnt2a* MO | ns | ns | p=0.0026 | p=0.0099 | p<0.001 | p<0.001 |
| 32 vs 36 hpf | *tnnt2a* MO | ns | ns | p=0.0026 | p=0.0026 | p=0.022 | p=0.031 |
| 28 hpf | *control* vs *tnnt2a* | ns | p=0.028 | p<0.001 | p<0.001 | p<0.001 | p<0.001 |
| 32 hpf | *control* vs *tnnt2a* | ns | ns | p<0.001 | p<0.001 | p<0.001 | p<0.001 |
| 36 hpf | *control* vs *tnnt2a* | ns | p=0.0035 | p<0.001 | p<0.001 | p<0.001 | p<0.001 |

**Supplementary Table 1** Statistical analysis related to Figure 1. For each vessel, differences between 1) control MO and *tnnt2a* MO groups and 2) time points were analyzed by Fisher's exact test. *p* values are listed within each cell, ns = not significant.

| Time of Collection | Embryo | Vessel Segment | | | | | |
|---|---|---|---|---|---|---|---|
| | | AA1 | ICA | CaDI | BCA | LDA | DA |
| 28 vs 32 hpf | SC | ns | ns | ns | ns | ns | ns |
| 28 vs 36 hpf | SC | ns | ns | ns | ns | ns | ns |
| 32 vs 36 hpf | SC | ns | ns | ns | ns | ns | ns |
| 28 vs 32 hpf | DM | ns | ns | ns | ns | p=0.01 | p<0.001 |
| 28 vs 36 hpf | DM | ns | ns | p=0.017 | p=0.008 | p<0.001 | p<0.001 |
| 32 vs 36 hpf | DM | ns | ns | ns (p=0.08) | ns (p=0.09) | p=0.002 | p=0.018 |
| 28 hpf | SC vs DM | ns | ns | p=0.0016 | p<0.001 | ns | ns |
| 32 hpf | SC vs DM | ns | ns | p=0.0075 | p<0.001 | ns | p=0.0075 |
| 36 hpf | SC vs DM | ns | p=0.0098 | p<0.001 | p<0.001 | p<0.001 | p<0.001 |

**Supplementary Table 2** Statistical analysis related to Figure 2. For each vessel, differences between 1) sibling control (SC) and double mutant (DM) groups and 2) time points were analyzed by Fisher's exact test. *p* values are listed within each cell, ns = not significant.

| Time of Collection | Embryo | Treatment | Vessel Segment | | | |
|---|---|---|---|---|---|---|
| | | | CaDI | BCA | LDA | DA |
| 36 hpf | SC | T vs C | p<0.001 | p<0.001 | p<0.001 | p<0.001 |
| 36 hpf | DM | T vs C | p=0.047 | p=0.008 | p=0.008 | ns |
| 40 hpf | SC | T vs C | ns | ns | ns | ns |
| 40 hpf | DM | T vs C | ns | ns | ns | ns |
| 36 vs 40 hpf | SC | C | ns | ns | ns | ns |
| 36 vs 40 hpf | SC | T | p<0.001 | p<0.001 | p<0.001 | p<0.001 |
| 36 vs 40 hpf | DM | C | ns | ns | ns | ns |
| 36 vs 40 hpf | DM | T | ns | ns | ns (p=0.056) | ns |
| 36 hpf | SC vs DM | C | p<0.001 | p<0.001 | p<0.001 | p<0.001 |
| 36 hpf | SC vs DM | T | ns | ns | ns | ns |
| 40 hpf | SC vs DM | C | p<0.001 | p<0.001 | p<0.001 | p<0.001 |
| 40 hpf | SC vs DM | T | p<0.001 | p<0.001 | p<0.001 | p<0.001 |

**Supplementary Table 3** Statistical analysis related to Figure 3. For each vessel, differences between 1) sibling control (SC) and *bmp10;bmp10-like* double mutant (DM) groups, 2) time points, and 3) treatments (tricaine: T, control: C) were analyzed by Fisher's exact test. *p* values are listed within each cell, ns = not significant.
